# Meta-analysis reveals reproducible rapamycin-induced shifts in the mouse gut microbiome

**DOI:** 10.1101/2025.09.12.675777

**Authors:** Asiri Ortega-Matienzo, Franziska Meiners, Patrick Newels, Ingmar Zude, Georg Fuellen, Israel Barrantes

**Affiliations:** Institute for Biostatistics and Informatics in Medicine and Ageing Research, Rostock University Medical Center, Rostock, Germany; Biovis Diagnostik, Limburg-Eschhofen, Germany

**Keywords:** rapamycin, ageing, gut microbiome, 16S rRNA, meta-analysis

## Abstract

Rapamycin is a geroprotective compound that extends lifespan and is well characterised in systemic physiology. However, due to heterogeneous methodologies, its effects on the gut microbiome, an emerging regulator of ageing and health, remain poorly defined. Here, we systematically assessed these effects in mice using a harmonised meta-analysis workflow, reprocessing raw sequence data end-to-end.

We conducted a harmonised meta-analysis of the gut microbiome using 16S rRNA gene amplicon data from three independent mouse studies (n = 54) spanning dietary and intraperitoneal rapamycin interventions with matched controls. All samples were uniformly processed using a standardised 16S rRNA bioinformatics workflow. Genus-level abundances were modelled using two complementary approaches: a per-study zero-inflated beta modelling framework with random-effects meta-analysis, and a batch-corrected linear modelling framework incorporating correction prior to meta-analysis. Dose-response meta-regression assessed whether rapamycin-associated changes scaled with dose (42–990 mg/kg), integrating presence and magnitude of effects across heterogeneous study designs.

Rapamycin treatment induced reproducible shifts in specific taxa across studies, with a subset of core responders consistently altered regardless of dose. Among the core responders, *Bacteroides* increased (OR = 1.59, q = 0.042) and *Muribaculum* decreased (OR = 0.55, q = 0.042), each significant in both frameworks. Whereas *Ruminococcus* decreased (OR = 0.60, q = 0.089), it was significant in the batch-corrected framework, but only a trend in zero-inflated modelling. Dose-sensitivity analyses highlighted Lachnospiraceae as the most consistent dose-sensitive taxon, with other associations largely restricted to extreme doses. Functional pathway analysis revealed alterations in energy metabolism and microbial stress response.

This harmonised meta-analysis highlights notable shifts in the gut microbiome connected to rapamycin across the examined studies, outlining potential core microbial signatures. These findings were observed within heterogeneous experimental contexts, and further validation in additional studies will be needed to confirm their generality.

## Introduction

Ageing is a primary risk factor for numerous diseases, including metabolic and cardiovascular disorders, cancer, and dementia, posing significant societal and economic challenges. Although life expectancy has increased, healthspan has not kept pace, making ageing the predominant driver for chronic illnesses (Guo et al., 2022). Understanding and modulating the fundamental biological ageing processes has become a central goal of geroscience, potentially enabling the simultaneous delay of multiple age-related diseases through interventions targeting core ageing mechanisms (Moqri et al., 2023).

The cumulative effects of cellular damage, impaired repair mechanisms, and alterations in various biological processes contribute to an increased vulnerability to chronic diseases as we age (Guo et al., 2022; Sprott, 2010). At the cellular and molecular level, ageing features interconnected changes in genome maintenance, proteostasis, mitochondrial quality control/autophagy, nutrient-sensing and metabolic pathways, cellular senescence, inflammation, and host–microbiome interactions (Zhao et al., 2022; Hipp et al., 2019; Lim et al., 2024; Saxton & Sabatini, 2017; McHugh & Gil, 2018; Ghosh et al., 2022).

Promising approaches to enhance healthy ageing include inhibition of nutrient-sensing pathways, clearance of senescent cells, exposure to systemic factors that rejuvenate stem-cell function, lifestyle interventions, and microbiome-targeted therapies such as prebiotics, probiotics, faecal microbiota transplantation and diet-based fibre/polyphenol interventions (Baker et al., 2016; Chenhuichen et al., 2022; Conboy et al., 2005; Kennedy & Lamming, 2016; Saxton & Sabatini, 2017; Yan et al., 2023; Meiners et al., 2025). Increasing autophagy, including mitophagy, and reducing age-associated inflammation have emerged as central mechanisms for these interventions (Partridge et al., 2020). Among the conserved pathways regulating ageing, the mTOR signalling network stands out as a promising therapeutic target, integrating nutrient, energy, and stress signals to coordinate cellular growth, metabolism, and survival (Papadopoli et al., 2019).

Rapamycin, a natural macrolide derived from bacteria, selectively inhibits mTORC1 and has been shown to extend lifespan and improve healthspan in diverse organisms, from yeast to mammals (Harrison et al., 2009; Powers et al., 2006; Jia et al., 2004; Kapahi et al., 2004; Fok et al., 2014). A recent comprehensive meta-analysis across 167 studies and eight vertebrate species confirmed the geroprotective efficacy of rapamycin, highlighting its comparable effectiveness to dietary restriction (Ivimey-Cook et al., 2025). In humans, rapamycin and related rapalogs are considered candidate gerotherapeutics, with small randomized trials in older adults showing improved antiviral immune signaling and a 2024 systematic review reporting benefits across immune, cardiovascular, and skin physiology, but lifespan and frailty endpoints remain untested (Mannick et al., 2021; Lee et al., 2024).

Besides mTOR as a potential target, the gut microbiome has emerged as both a modulator and biomarker of healthy ageing, with age-related compositional changes characterised by reduced microbial diversity, loss of beneficial taxa, and increased pro-inflammatory bacteria which strongly correlate with frailty, chronic inflammation, metabolic disorders, and immune dysregulation in older individuals (Biagi et al., 2016; Ravikrishnan et al., 2024; Han et al., 2017; Lau et al., 2021; Wilmanski et al., 2021; Nagpal et al., 2018). Gut microbial metabolites such as short-chain fatty acids (SCFAs) directly affect host metabolism, inflammation, and immunity, making the microbiome a key axis in ageing biology, as shown by dietary interventions (Meiners et al., 2025).

Consistent with its roles in nutrient sensing, inflammation and metabolism, rapamycin has been reported to modulate the gut microbiome. Across animal studies, the direction and magnitude of these changes vary with dose, route, treatment duration, age, and disease context. In mice, dietary or encapsulated rapamycin in middle-to late-life induced modest but reproducible shifts, including *Bacteroides* enrichment and segmented filamentous bacteria increases, often persisting after treatment cessation (Hurez et al., 2015; Bitto et al., 2016). By contrast, chronic intraperitoneal regimens in young mice induced stronger community reweighting, marked by reduced diversity, distinct separation, and depletion of *Akkermansia*, coinciding with adverse metabolic phenotypes (Han et al., 2021).

A recent time-course study showed delayed onset of these effects, with diversity and functional changes only emerging after 30 days (Wu et al., 2025). Rapamycin improved barrier integrity in inflammatory models and reduced inflammation, often accompanied by *Lactobacillus* increases and Proteobacteria decreases (Guo et al., 2023; Xu et al., 2020; Zhang et al., 2020). Beyond mice, rat and early human studies similarly report shifts in butyrate-associated taxa and SCFA profiles (Bhat et al., 2017; Lin et al., 2025 preprint). Given these context and method differences, the direction and magnitude of rapamycin’s microbiome effects remain heterogeneous and not directly comparable, leaving shared signatures unresolved.

To address this gap, we uniformly reprocessed mouse 16S rRNA datasets and synthesised results across studies within a harmonised analysis workflow. Our objectives were to (1) identify reproducible microbial signatures of rapamycin treatment across independent studies, (2) assess dose-response relationships between rapamycin exposure and microbiome composition, and (3) evaluate the consistency of these effects across different experimental conditions.

Our meta-analysis of 54 samples across three independent studies revealed that rapamycin is associated with consistent genus-level shifts in gut microbiome composition, with significant effects on key genera, including *Bacteroides*, *Muribaculum*, and *Ruminococcus* as well as taxon-specific dose sensitivity. These results suggest common microbiome responses to rapamycin in the contexts studied, although further independent studies are needed to determine how broadly these patterns apply.

## Methods

### Study Selection and Data Sources

We conducted a meta-analysis of 16S rRNA amplicon sequencing data from three independent mouse gut microbiota studies: Bitto2016 (PRJEB13256) — Middle-aged C57BL/6J mice received rapamycin either by intraperitoneal (IP) injection (8.0 mg/kg daily for 3 months) or dietary supplementation (126 ppm for 90 days), with matched controls (Bitto et al., 2016). Yang2022 (PRJNA717340) — wild-type male C57BL/6J mice were treated with short-term IP rapamycin (17.5 mg/kg daily for 14 days) in a dextran sodium sulfate (DSS) colitis model; only wild-type treatment and control groups were included in this analysis (Yang et al., 2022). Han2021 (PRJNA436876) — Young male C57BL/6 mice received long-term IP sirolimus (rapamycin; 0.5 mg/kg daily for 12 weeks) with or without probiotic supplementation; the probiotic arm was excluded to maintain intervention consistency (Han et al., 2021). The Han2021 dataset involved DSS-colitis induction; although we analysed only wild-type animals, this inflammatory context may modulate responses. Effects should therefore be read as context-averaged. All datasets were obtained from the European Nucleotide Archive (ENA) and NCBI Sequence Read Archive (SRA) (Harrison et al., 2020; Leinonen et al., 2011) and are summarised in Table 1.

**Table 1.**
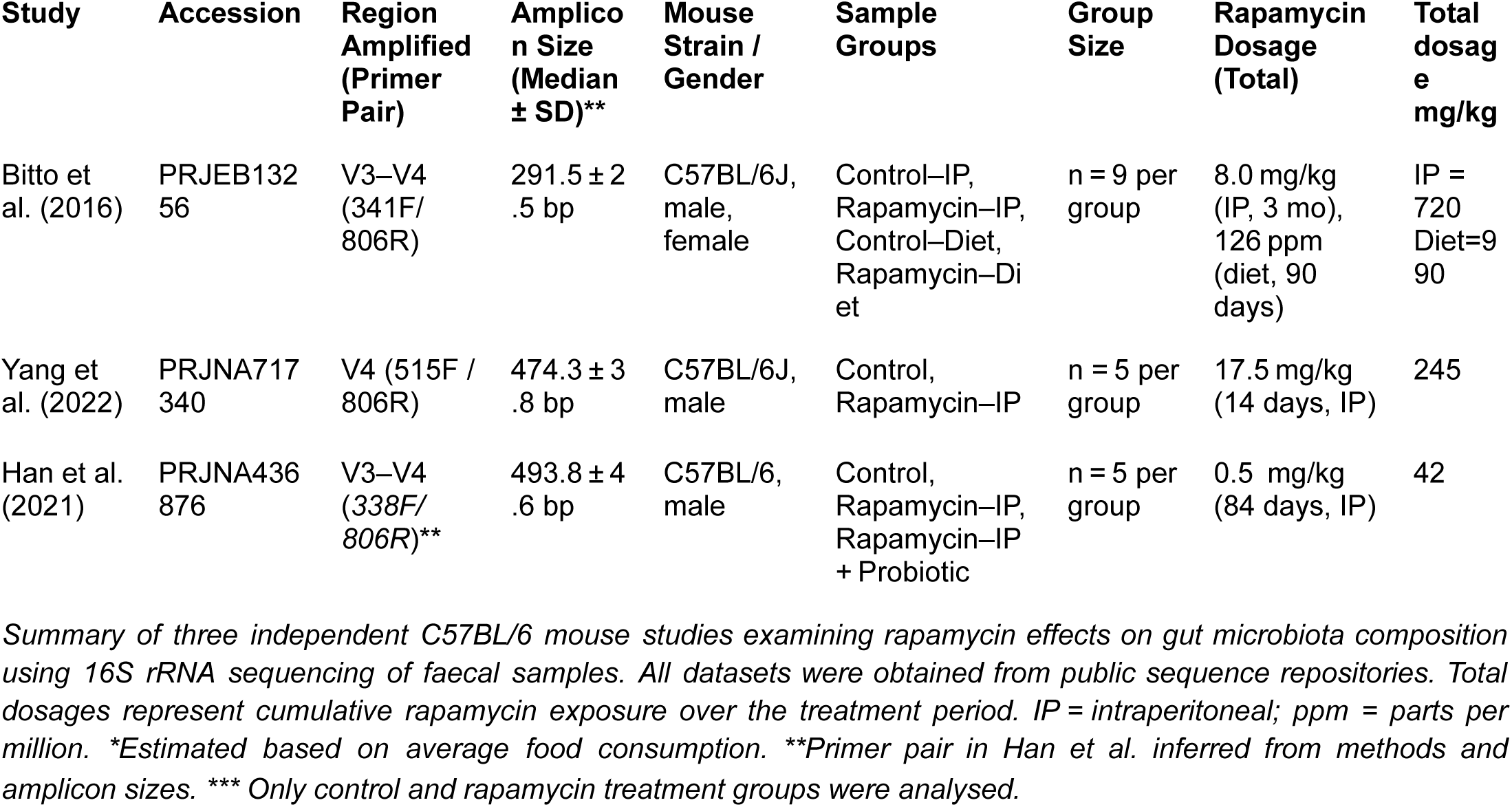
Characteristics of studies included in rapamycin gut microbiome meta-analysis.

Studies met the following inclusion criteria: (1) C57BL/6 mice as host organism, (2) faecal samples as source material, (3) Illumina sequencing platform, (4) rapamycin treatment versus control comparison, and (5) availability of raw sequencing data and metadata. Except for these three, no other studies matched these criteria.

### Sequence preprocessing and taxonomic classification

Raw sequencing data (FASTQ files) were processed using QIIME2 v2020.8.0 (Bolyen et al., 2019) within a Singularity v3.8.7 container environment. Data import used the PairedEndFastqManifestPhred33V2 format, followed by quality control visualisation of demultiplexed sequences, and denoising using the DADA2 algorithm (Callahan et al., 2016), which generates amplicon sequence variants (ASVs) by explicitly modelling and correcting sequencing errors. We chose to retain ASVs rather than cluster into operational taxonomic units (OTUs) because ASVs provide higher-resolution, reproducible sequence labels that are directly comparable across studies and datasets, avoiding the loss of biological information inherent in fixed-threshold clustering, which achieves close agreement with theoretical community composition (Abellan-Schneyder et al., 2021).

For each dataset, DADA2 truncation parameters were optimised based on quality profiles: Bitto2016 and Yang2022 were processed with TRUNC_LEN_F = 250, TRUNC_LEN_R = 250, while the Han2021 dataset required a longer forward read truncation (TRUNC_LEN_F = 270) to preserve high-quality bases. All datasets were processed without primer trimming to facilitate consistent cross-study analysis despite using different primer sets (Bitto2016, Han2021: V3–V4; Yang2022: V4).

Taxonomic classification employed the SILVA 138 99% OTUs full-length sequences classifier (Quast et al., 2013) at a confidence threshold of 0.7. Downstream quality control included removing chloroplast and mitochondrial sequences, excluding non-bacterial sequences and ASVs lacking phylum-level classification, and filtering samples with fewer than 2,000 reads. Phylogenetic trees were built using FastTree v2.1.10 (Price et al., 2010).

Functional pathways were predicted using PICRUSt2 v2.5.3 (Douglas et al., 2020) with default parameters (NSTI ≤ 2, minimum read count = 1, alignment fraction ≥ 0.8). Stratified outputs were retained to enable taxon-specific pathway contribution analysis, and copy-number normalisation was applied as recommended.

### Re-analysis of individual studies

We re-analysed the microbiome data using the exported QIIME2 outputs, which included feature tables, taxonomy, phylogenetic trees and metadata. All analyses were performed in R v4.4.2 with several key packages: phyloseq v1.50.0 (McMurdie & Holmes, 2013) for microbiome data handling, DESeq2 v1.46.0 for differential abundance testing (Love et al., 2014), microbiome v1.28.0 for core microbiome analysis (Lahti et al., 2017), and vegan v2.6-8 for ecological diversity metrics (Oksanen et al., 2018). The data analysis workflow began by creating phyloseq objects that combined all exported features. We applied quality control filters to remove low-quality samples (<2,000 reads), non-bacterial sequences, and ASVs without phylum-level classification.

Taxonomic composition was assessed with detection thresholds of 1.25% at genus level and 0.1% at phylum level (≥50% prevalence), consistent with thresholds shown to capture core genera across ≥80% of subjects in diverse cohorts (Shetty et al., 2017). Firmicutes-to-Bacteroidetes (F/B) ratios were calculated for each treatment group and compared using Wilcoxon rank-sum tests (p < 0.05). Core microbiome taxa were defined as present at ≥ 0.1% abundance in ≥ 80% of samples per group and compared using binary presence/absence matrices.

Alpha diversity was calculated using the Shannon diversity index. Between-group differences in alpha diversity were assessed using Wilcoxon rank-sum tests for pairwise comparisons and Kruskal-Wallis tests when more than two groups were present. Beta diversity was determined using weighted and unweighted UniFrac distances and Bray-Curtis dissimilarity. For Bray-Curtis analysis, we calculated the divergence of each sample from its group median to assess within-group variation, following Salonen et al. (2014). Between-group differences in beta diversity were tested using PERMANOVA with 10,000 permutations. Ordination was performed using Principal Coordinates Analysis (PCoA) to visualise clustering patterns between treatment groups.

Differential abundance analysis was performed using DESeq2 with Benjamini-Hochberg false discovery rate (FDR) correction for multiple testing. For the Bitto2016 dataset, a relative abundance filtering threshold of 0.1% was applied; for the Yang2022 and Han2021 datasets, a more stringent 0.25% threshold was used to account for differences in sequencing depth. Taxa with adjusted p-values < 0.05 and |log2FoldChange| > 3 were considered significantly differentially abundant.

Pathway abundance tables were filtered to include only pathways with a minimum count of 1,000 across all samples. Differential pathway abundance analysis was performed using DESeq2, with statistically significant pathways defined at an adjusted p-value < 0.05 after Benjamini-Hochberg correction for multiple testing. Log^2^ fold change values were calculated for each pathway, and pathway identifiers were mapped to the MetaCyc database (Caspi et al., 2020) to facilitate biological interpretation.

### Meta-Analysis Framework

To address methodological heterogeneity and facilitate cross-study comparisons, we implemented two complementary meta-analytical frameworks: an MMUPHin-based approach incorporating batch correction to harmonise datasets prior to meta-analysis, and a GAMLSS + metamicrobiomeR framework that models each study individually, followed by random-effects meta-analysis. Using both frameworks allowed us to assess the sensitivity of findings to different modelling assumptions (batch-corrected linear modelling versus zero-inflated beta modelling) and to report method-specific results transparently. We present results from both, highlighting concordant signals and clearly indicating method-specific findings, so that conclusions do not hinge on a single analytic choice.

We standardised sample metadata and treatment coding across all datasets for valid cross-study comparisons. To maximise sample size, we excluded sex and age as covariates despite their availability in most datasets. Consequently, potential sex and age effects cannot be distinguished from study-specific effects in our meta-analysis. Uniform variables were generated for primary treatment (“StandardTreatment”: Control/Rapamycin), administration route (“AdminRoute”: Food/IP), and cumulative rapamycin exposure (daily and total dose in mg/kg, with log-transformed versions for regression modelling). Standardised cumulative doses were: Han2021 (42 mg/kg), Yang2022 (245 mg/kg), Bitto2016-IP (720 mg/kg), and Bitto2016-Diet (990 mg/kg). Previously processed genus-level abundance tables were further harmonised by standardising taxon names and removing all “unclassified,” “uncultured,” or “unknown” genera, providing a consistent starting point for both analytical frameworks.

### Batch correction and cross-study harmonisation

To address the technical and methodological differences between studies that could mask biological signals across studies, we applied batch correction using the Meta-Analysis Methods with a Uniform Pipeline for Heterogeneity in microbiome studies (MMUPHin) package v1.14.0 (Ma et al., 2022). This method harmonises microbiome data across heterogeneous studies by adapting the ComBat algorithm (Johnson et al., 2007) for zero-inflated, sparse datasets. The merged genus-level abundance table was Total-Sum Scaled (TSS), filtered for prevalence (≥ 1 × 10^-5^ in ≥ 2 samples), and batch-corrected, including administration route (‘AdminRoute’) as a covariate. We did not include age or sex as a covariate, because these variables were constant or incomplete within individual studies and were highly collinear with study. In ComBat-style models, covariates without within-batch variation are not identifiable and can induce over-correction. Effectiveness was evaluated via Principal Coordinates Analysis (PCoA) and PERMANOVA.

For meta-analytical differential abundance testing, we used the lm_meta function from MMUPHin on Arcsin Square Root (AST) transformed data, applying Linear Models (Mallick et al., 2021) with random-effects meta-analysis (Viechtbauer, 2010). MaAsLin2 models were run separately per study with ‘StandardTreatment’ as exposure and ‘AdminRoute’ as a covariate. Study-specific effect sizes and standard errors were pooled using the MMUPHin random effects framework, with heterogeneity quantified by I², τ², and the Cochran Q statistic.

As a complementary analysis, we examined only rapamycin-treated samples (n = 27) fitting genus-specific linear models with standardised log-transformed total dose as predictor and study as covariate. This within-treatment approach reduced control–treatment contrast to detect subtle dose gradients across the 23-fold range (42–720 mg/kg). Significance was determined by Benjamini-Hochberg correction (q < 0.05).

### Zero-inflated beta regression meta-analysis

Following taxonomic harmonisation, genus-level abundance data were converted to relative proportions (0–1 scale) to preserve the natural zero structure of the data. We modelled abundances for each study independently using Generalised Additive Models for Location, Scale and Shape (GAMLSS; gamlss v5.4-22) with zero-inflated beta regression (BEZI family), which appropriately handles both the compositional nature of microbiome data and the high frequency of zeros commonly observed in these datasets (Stasinopoulos et al., 2012). Zero-inflated beta models were fitted for each genus with StandardTreatment as a binary factor contrasting control versus rapamycin treatment, allowing us to capture both the probability of a genus being absent (zero component) and its relative abundance when present (beta component).

Genera with SE > 5 were excluded to avoid unstable estimates. Only taxa present in ≥ 2 studies were retained for meta-analysis. Study-specific effect sizes (BEZI μ logit coefficients) and standard errors were combined using random-effects meta-analysis via metamicrobiomeR v1.2 (Ho et al., 2018). Results are reported as odds ratios (OR) with 95% confidence intervals, where OR > 1 indicates increased abundance and OR < 1 indicates decreased abundance. Statistical significance was set at FDR q < 0.05, with heterogeneity assessed using Cochran’s Q test and the DerSimonian-Laird estimator of τ² and I².

We conducted meta-regression using total cumulative dose as a continuous moderator to investigate dose-response relationships. The Bitto2016 study was subdivided by administration route (intraperitoneal vs. dietary) to account for distinct pharmacokinetic profiles and dosing regimens, resulting in four study arms for dose-response modelling. Meta-regression models were fitted using metafor v4.8-0:

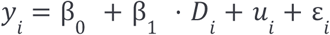

Where 𝑦*_i_* represents the log OR from GAMLSS analysis, β_1_ represents the dose-response. 𝐷*_i_* is the total cumulative rapamycin dose (mg/kg) for study arm 𝑖 slope, 𝑢*_i_* represents random study effects, and ε*_i_* represents sampling error. Significance was assessed via Wald tests with Benjamini-Hochberg FDR correction. Sensitivity analyses excluded the highest-dose arm (Bitto2016-Diet), restricting the range to 42–720 mg/kg. Taxa were analysed within any subset only if ≥ 3 study arms had estimable effects.

### Visualisation

Results visualisation included heatmaps showing significant taxa effect size patterns across studies, PCoA plots displaying community structure before and after batch correction, forest plots displaying individual study and pooled meta-analysis results, treatment effect plots, and dose-response analysis plots. Heatmaps used the metamicrobiomeR v1.2 visualisation functions, while forest plots and dose-response plots employed custom ggplot2 v3.5.2 implementations. Multiple plots were combined using patchwork v1.3.0 (Pedersen, 2022). Statistical significance was set at FDR-adjusted p < 0.05 for primary analyses and p < 0.10 for exploratory visualisations.

**Fig. 1.**
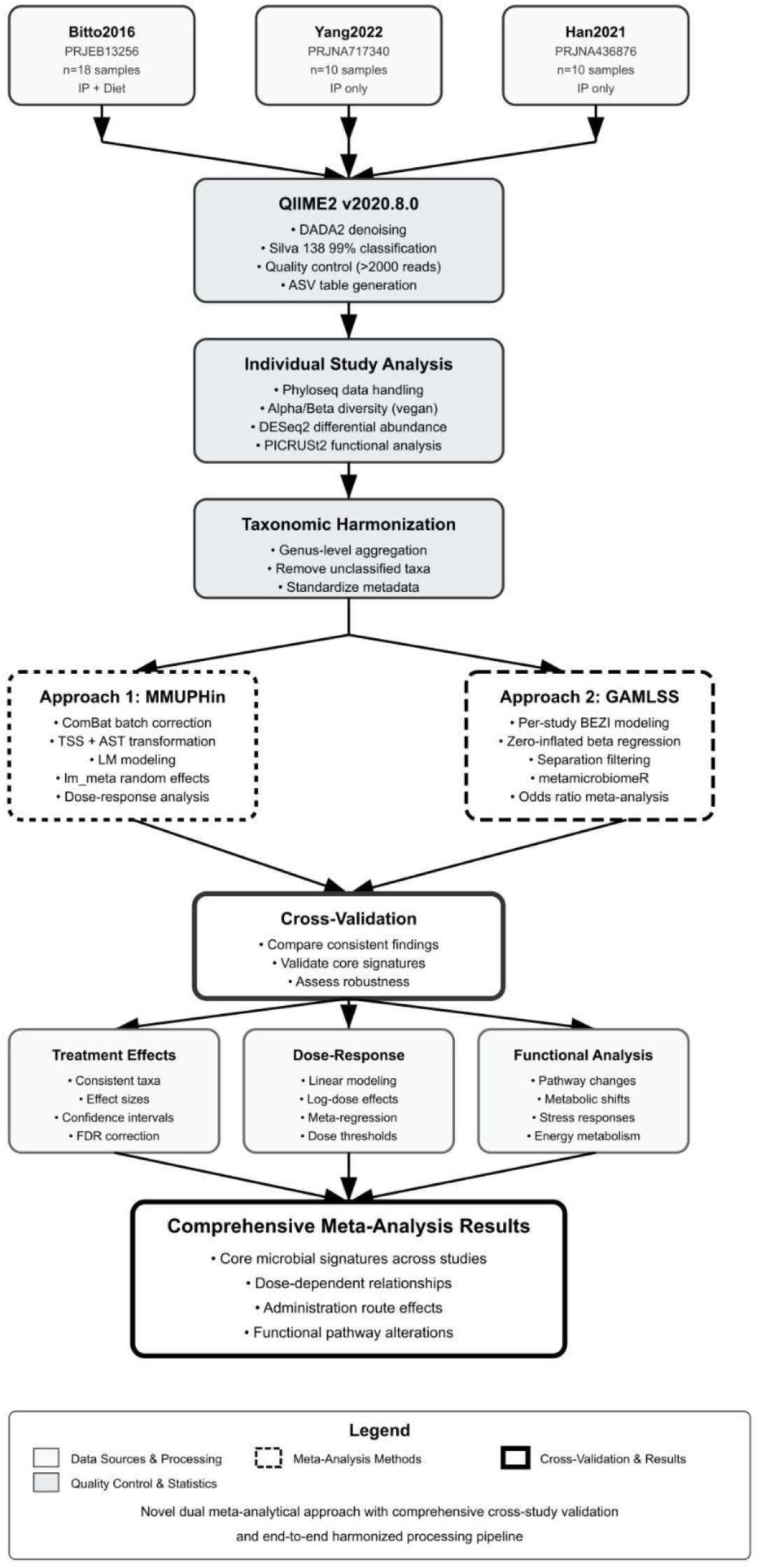
Harmonised meta-analysis pipeline for rapamycin-microbiome studies: dual MMUPHin and GAMLSS approaches with cross-study validation

## Results

### Single study analyses

#### Administration routes modulate both taxonomic and metabolic microbiome responses

We analysed each dataset individually and then combined them to provide a comprehensive understanding of the effect of rapamycin on the microbiome. Bitto et al. 2016 investigated whether transient rapamycin treatment could improve lifespan and healthspan in middle-aged mice, specifically examining whether short-term administration could benefit ageing-related processes. They utilised 19–20-month-old C57BL/6J mice, which were divided into two treatment groups: intraperitoneal (IP) injection (8.0 mg/kg daily) and dietary supplementation (126 ppm), both administered for 90 days. Faecal samples were collected per cage at 3 months of treatment, with each sample representing pooled material from multiple co-housed mice.

Upon uniform taxonomic reassignment, we found route-dependent responses to rapamycin treatment in the Bitto2016 dataset, representing moderate doses (720 mg/kg IP, 990 mg/kg food) administered to aged mice (20–21 months) over 90 days. Taxonomic composition showed Firmicutes and Bacteroidetes dominance with no significant phylum-level changes (p = 0.717). The F/B ratio significantly decreased with IP rapamycin (0.67 vs 0.89, p = 0.031) but showed only trends with dietary administration (0.83 vs 1.35, p = 0.13).

Alpha diversity remained unchanged across both administration routes (Fig. S14). In contrast, beta diversity revealed significant community restructuring (PERMANOVA p < 0.0001 for both weighted/unweighted UniFrac) (Fig. S17, S20), indicating significant differences in both microbial composition and their relative abundances between administration routes, with samples clustering primarily by administration route rather than treatment, highlighting route-specific microbiome modulation mechanisms.

Differential abundance analysis revealed similar effects between routes, with one key distinction. Both IP and dietary administration significantly enriched members of the Muribaculaceae family, with IP showing uniform enrichment (LFC > 20) alongside *Lacrimispora sp*. (LFC > 23). In contrast, dietary rapamycin uniquely showed complex bidirectional effects on Muribaculaceae ASVs (LFC > 20/ LFC > -20) alongside depletion of several taxonomically unresolved bacterial ASVs (*Unclassified/Uncultured* groups, LFC > -20)(Fig. S25).

Functional pathway analysis showed distinct metabolic modulation patterns between routes. IP treatment upregulated sucrose biosynthesis pathways (Log_2_FC > 10) and showed a modest downregulation in amino acid biosynthesis. In contrast, dietary administration affected osmotic regulation through glycine betaine degradation (Log_2_FC > 10) and altered cell wall biosynthesis via peptidoglycan pathways (Log_2_FC > 10). These findings demonstrate that the administration route modulates both taxonomic and metabolic microbiome responses, despite both routes targeting fibre-fermenting bacteria (Fig. S28).

#### Moderate rapamycin dose increases diversity while preserving community structure

Yang et al. investigated whether RhoB-driven autophagy shapes colitis and the microbiome. Male C57BL/6J mice received short-term intraperitoneal rapamycin (17.5 mg/kg/day, 14 days) in a DSS model. For our reanalysis, we restricted to the wild-type cohort and contrasted rapamycin vs vehicle to estimate rapamycin-specific microbiome effects independent of RhoB genotype.

The overall microbial community structure remained stable at the phylum level, although subtle shifts emerged, as seen in our analysis using a harmonised workflow. Upon examination of taxonomic composition, Firmicutes and Bacteroidetes dominance was maintained with no significant phylum-level alterations (p = 0.462) and minimal F/B ratio change (1.16 vs 1.11, p = 0.84) (Fig. S8).

Alpha diversity significantly increased with rapamycin treatment (p = 0.032). Beta diversity showed no significant community-level changes in weighted UniFrac (PERMANOVA p = 0.39) but revealed altered taxon presence/absence patterns in unweighted UniFrac (dispersion p = 0.039). This indicates changes in microbial composition rather than abundance, with minimal Bray-Curtis effects (p = 1.0). These findings suggest subtle taxonomic restructuring without gross community shifts (Fig. S15).

Differential abundance analysis revealed significant taxonomic changes following rapamycin treatment. Rapamycin increased the abundance of members of the Prevotellaceae family, *Alloprevotella*, *Mucispirillum* (Deferribacterota phylum), *Helicobacter*, and several Firmicutes members. Conversely, multiple Bacteroidetes representatives, *Akkermansia*, *Lactobacillus*, and Clostridia *UCG-014* showed significant decreases. *Muribaculum*, *Alistipes* and members of the Muribaculaceae family exhibited complex responses, with significantly increased and decreased abundances within these taxonomic groups (Fig. S26).

Functional pathway analysis revealed extensive metabolic modulation with upregulation of aromatic amino acid biosynthesis (chorismate), stress adaptation pathways (PPGPPMET-PWY), multiple energy metabolism variants (TCA cycles, glycolysis, butanediol biosynthesis), and oxygen-independent heme biosynthesis, collectively indicating that rapamycin triggers comprehensive metabolic adaptation with preserved community structure but fundamental functional reorganisation (Fig. S29).

#### Low-dose rapamycin reduced microbiome diversity

Within the single-study reanalysis, we next examined Han2021, a low-dose, long-term intraperitoneal rapamycin exposure. Han et al. (2021) investigated the metabolic effects of long-term rapamycin treatment and the role of the intestinal microbiota. They utilised young male C57BL/6 mice at five weeks old, treating them with IP sirolimus (rapamycin) at 0.5 mg/kg daily for 12 weeks. The study examined how chronic low-dose rapamycin affects metabolic parameters and whether associated microbiome changes contribute to observed metabolic disorders.

The dataset represented the lowest dose study (42 mg/kg) over 90 days. Our analysis using the harmonized workflow showed no significant phylum-level shift overall (p = 0.885), though *Alloprevotella* emerged as the most abundant genus, distinguishing the baseline of this study community structure from others where *Alistipes* (Bitto2016) and *Lactobacillus* (Yang2022) dominated. The F/B ratio significantly decreased with rapamycin treatment (p = 0.016), representing the strongest and most consistent phylum-level effect observed across all studies.

Core microbiome analysis showed that both control and rapamycin groups shared common taxa (members of the *Muribaculaceae* family, *Alloprevotella*, Lachnospiraceae NK4A136). However, rapamycin-treated groups lost *Akkermansia*, *Blautia*, and multiple *Eubacterium* species that remained present in controls, while gaining *Helicobacter* as a core member, a pattern observed across two of three studies.

Alpha diversity significantly decreased with rapamycin treatment (p = 0.016), contrasting sharply with the increased diversity observed in moderately dosed Yang2022 (p = 0.032) and unchanged diversity in high-dosed Bitto2016. Beta diversity analysis revealed the most pronounced treatment effects with significant compositional differences in both weighted UniFrac (PERMANOVA p = 0.0004) and unweighted UniFrac (p = 0.007) analyses, clear separation along the first principal coordinate in PCoA plots, and extreme data sparsity (93% zeros) likely contributing to Bray-Curtis dissimilarity values of 1.0 for all samples, indicating profound community wide ecological disruption rather than the subtle restructuring observed in lower-dose studies.

Differential abundance analysis demonstrated complex bidirectional patterns with rapamycin treatment significantly enriching several taxa including *Helicobacter*, *Bacteroides*, *Coriobacteriaceae* UCG-002, *Clostridibacter*, and Prevotellaceae UCG-001, while depleting others, including Prevotellaceae *NK3B31*, *Eubacterium xylanophilum* group, *Oscillibacter*, and *Alistipes*. Bacterial taxa within the same taxonomic groups of genera (*Alloprevotella*, Lachnospiraceae NK4A136) and family (Muribaculaceae) showed opposite responses to rapamycin treatment. In contrast, some became enriched, others decreased, indicating heterogeneous responses rather than uniform genus- or family-level effects.

The observed changes demonstrate that low-dose rapamycin substantially alters gut microbiome composition and diversity. However, differential pathway analysis would be needed to determine the biological significance of these changes for host health.

### Batch-corrected abundance meta-analysis

To quantify treatment effects after removing study structure, we performed a batch-corrected between-group meta-analysis. Following quality control and taxonomic harmonisation, the harmonised dataset comprised 54 samples and 112 genera across three studies, with 62 genera (55.4%) representing a conserved core microbiome (Fig. 2).

**Fig. 2.**
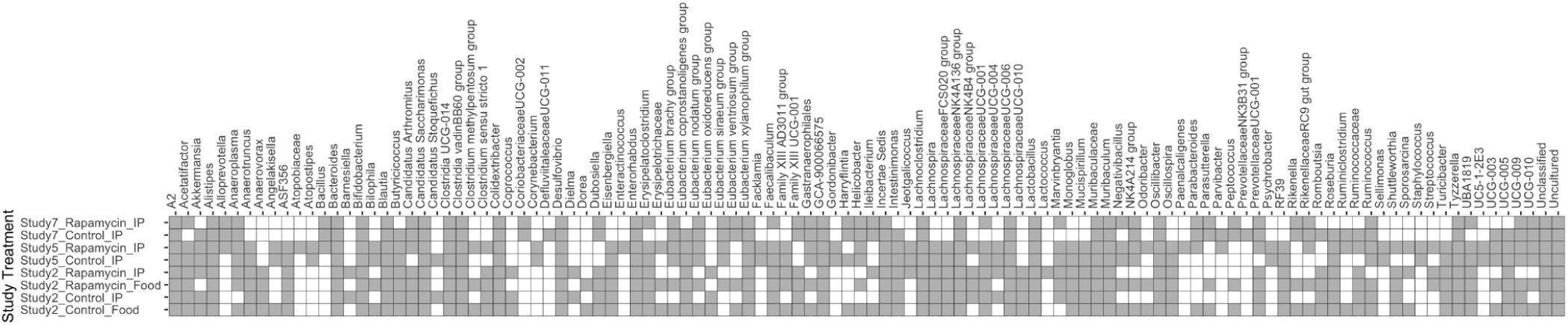
Presence-absence heatmap of gut microbial genera across rapamycin studies. Gray cells indicate that a given genus was detected (relative abundance > 0) in that treatment arm and white cells indicate it was not detected. Rows represent eight treatment arms across groups across Bitto2016, Yang2022 and Han2021 (control vs rapamycin; intraperitoneal (IP) vs dietary) and columns show the 112 genera passing prevalence and abundance filters. This visualisation highlights both the core taxa consistently present across studies and the study- or route-specific genera that may underlie differential dose-response signatures.

Initial PCoA of uncorrected data revealed strong study-specific clustering, with the first two principal coordinates explaining 33.1% and 28.3% of the variance. PERMANOVA confirmed a significant study effect (R² = 0.437, p < 0.001), exceeding the treatment effect (R² = 0.028, p = 0.168). Batch correction reduced study-specific clustering, with the first two coordinates explaining 31.1% and 19.5% of variance, and PERMANOVA showing decreased study effect (R² = 0.168, p < 0.001) alongside increased treatment effect (R² = 0.047, p = 0.015), consistent with removal of technical structure and enhancement of the biological signal (Fig. 3).

**Fig. 3.**
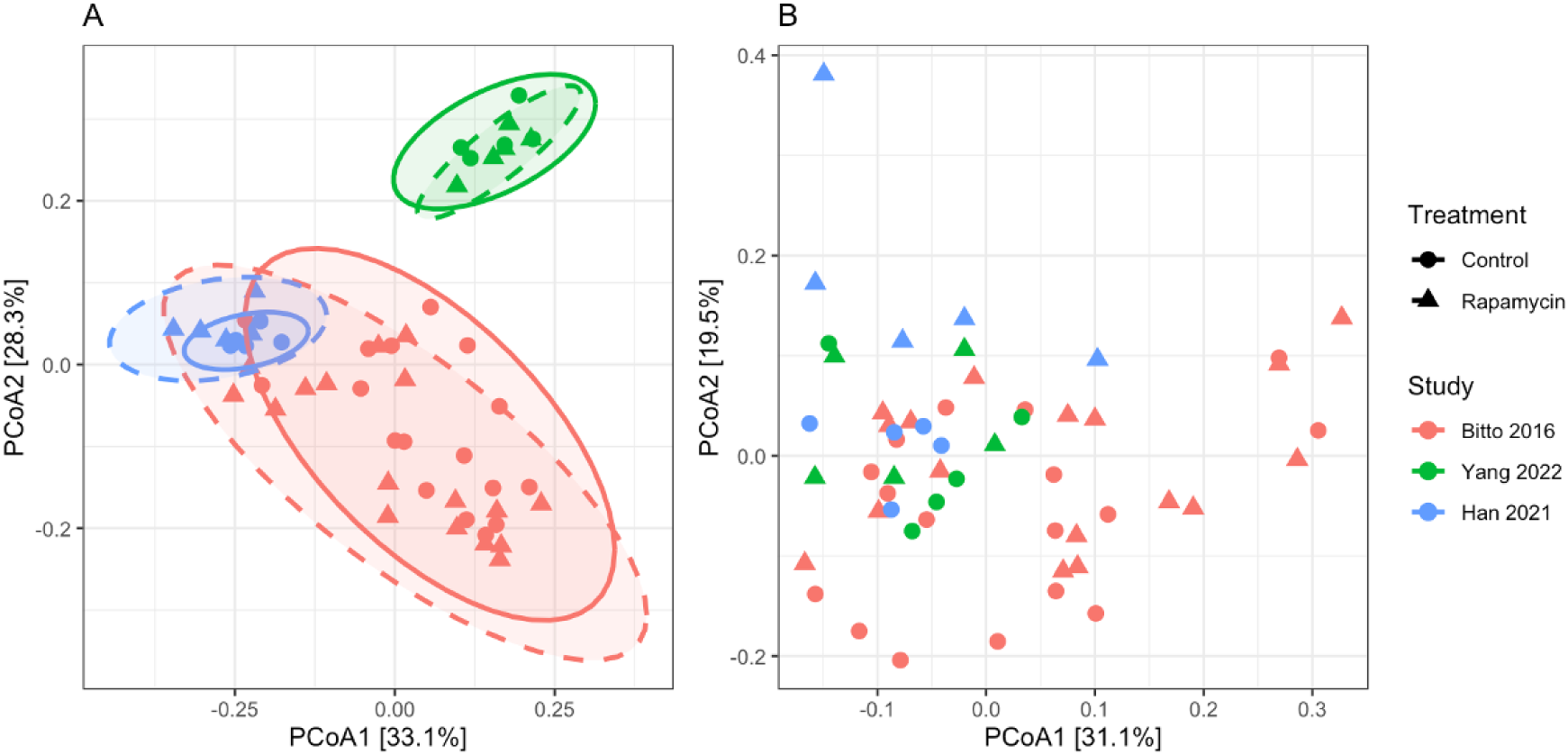
Effect of batch correction on community structure. Principal Coordinates Analysis (PCoA) based on Bray-Curtis dissimilarity before (A) and after (B) MMUPHin batch correction. (A) Before correction shows pronounced study-specific clustering with three distinct groups corresponding to Bitto 2016 (red points, lower region), Yang 2022 (green points, upper right), and Han 2021 (blue points, upper left). Ellipses indicate 95% confidence regions for study-treatment combinations, with solid and dashed lines distinguishing control (circles) and rapamycin (triangles) treatments within each study. The first two axes capture 61.4% of total variance (PCoA1: 33.1%, PCoA2: 28.3%). (B) After batch correction, samples from different studies are intermixed across the ordination space, with the first two axes explaining 50.6% of variance (PCoA1: 31.1%, PCoA2: 19.5%). Batch correction reduced study-specific clustering (R²: 0.44 to 0.16) and increased treatment effect detection (R²: 0.03 to 0.05).

Despite minimal community-level effects, meta-analysis identified four genera with statistically significant and consistent responses to rapamycin across studies (FDR < 0.05). *Bacteroides* showed the largest positive association (coefficient = +0.061 ± 0.016, q = 0.007), increasing in all three datasets. ASF356 was also enriched (coefficient = +0.025 ± 0.007, q = 0.007) in the two studies where it was present (absent in Han2021). In contrast, two fibre-degrading taxa were consistently depleted: *Ruminococcus* (coefficient = −0.023 ± 0.007, q= 0.023) and *Muribaculum* (coefficient = −0.032 ± 0.010, q = 0.023). A side-by-side summary of these MMUPHin results with the GAMLSS estimates is provided in Table 2.

**Table 2.**
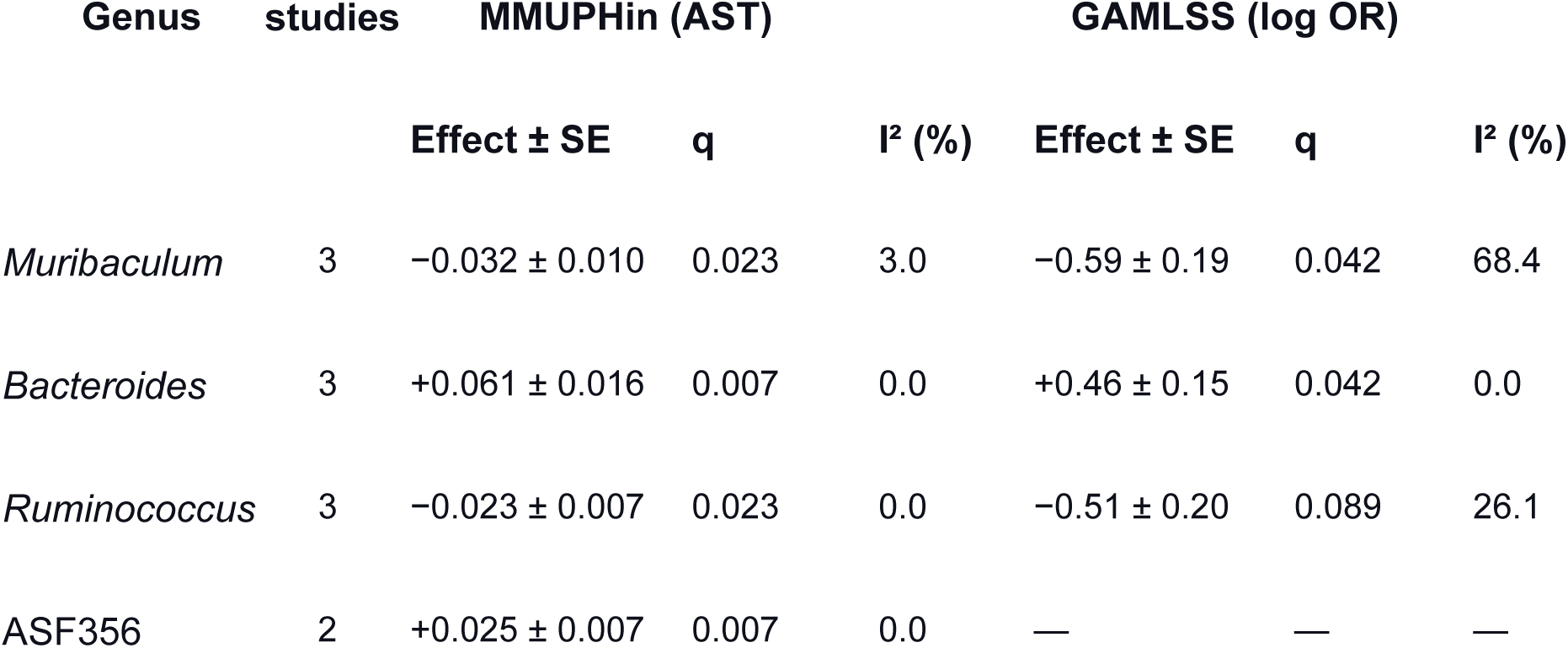
Significant genera across methods.

Cross-study consistency was high, with *Bacteroides*, *Ruminococcus*, and ASF356 showing no heterogeneity across studies (I² = 0%), while *Muribaculum* showed minimal heterogeneity (I² = 3%) (Fig. 4; Table 2).

**Fig. 4.**
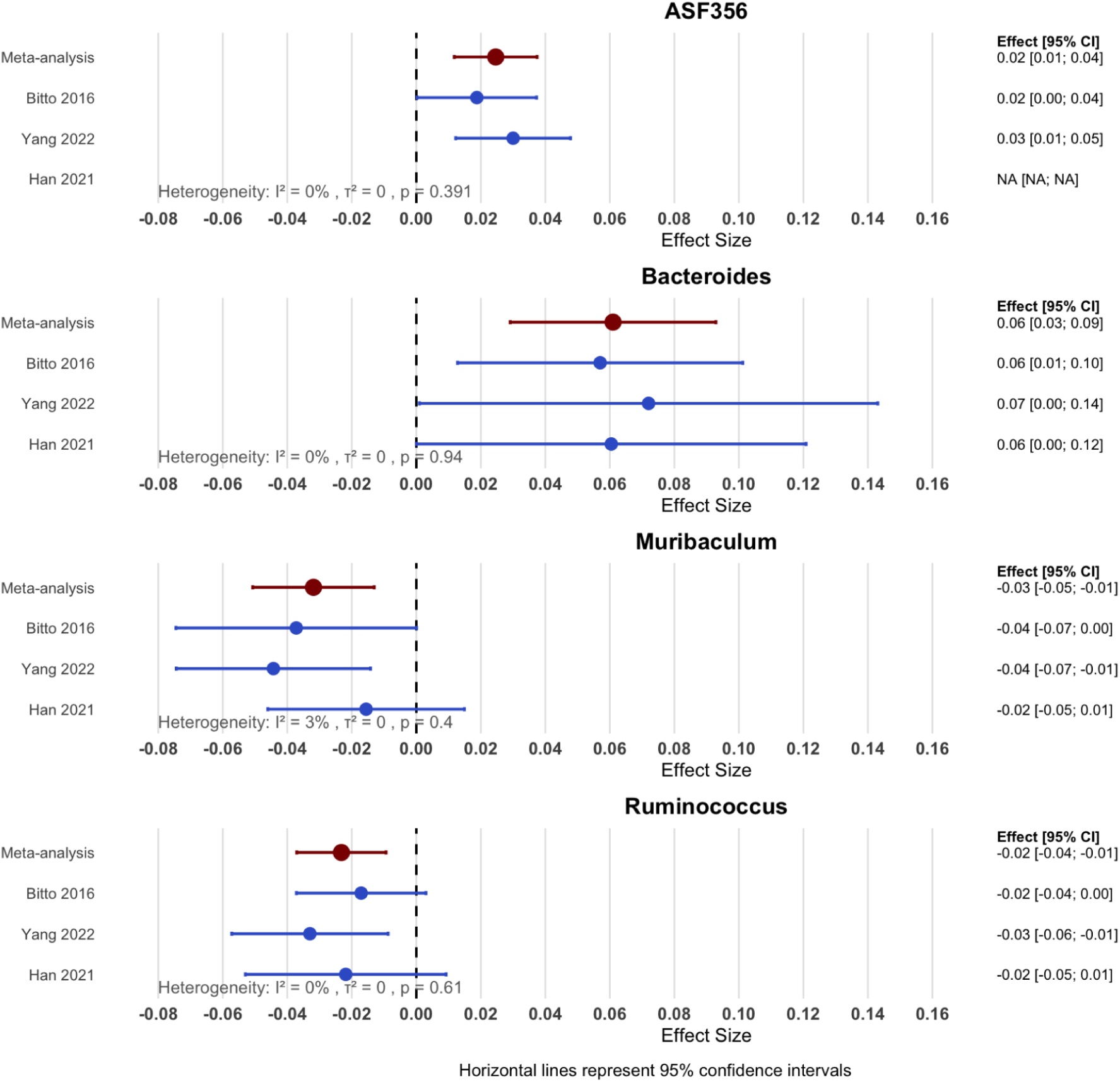
Forest plot of rapamycin effects on gut bacterial genera. Meta-analysis using random effects (RE) models pooled results across studies to identify four genera with significant treatment effects (FDR < 0.05). *Bacteroides* showed the most substantial enrichment with rapamycin treatment (+0.061 ± 0.016, q = 0.007, k = 3 studies), while *ASF356* was also significantly enriched (+0.025 ± 0.007, q = 0.007, k = 2 studies). Conversely, *Ruminococcus* and *Muribaculum* were significantly depleted by rapamycin treatment (*Ruminococcus*: -0.023 ± 0.007, FDR = 0.023; *Muribaculum*: -0.032 ± 0.010, q = 0.023). All four genera analysed here showed directionally consistent effects with low observed heterogeneity (I² ≤ 3%) within the included studies, indicating that these treatment-associated differences were stable across the analysed datasets despite differences in dosing regimens and administration routes. Study weights were determined by inverse variance, with differential contributions reflecting sample sizes and measurement precision. Meta-analysis was performed using the lm_meta function of MMUPHin.

### Dose-Response Analysis

Cross-study dose-response analysis using linear models, restricted to rapamycin-treated samples (n = 27) and adjusting for study, identified six genera with significant dose-related associations (FDR < 0.05). *Alistipes* showed the strongest decrease in relative abundance with dose (β = −0.384 ± 0.055, q = 5.05 × 10^-5^). Additional dose-related decreases in relative abundance were observed in the genera *Bacteroides*, Lachnospiraceae UCG-010 and UCG-005, whereas *Negativibacillus* and *Eubacterium nodatum* group showed dose-related increases in relative abundance. No significant association showed no detectable between-study heterogeneity (I² = 0%), indicating consistent dose-response patterns across studies (Fig. 5).

**Fig. 5.**
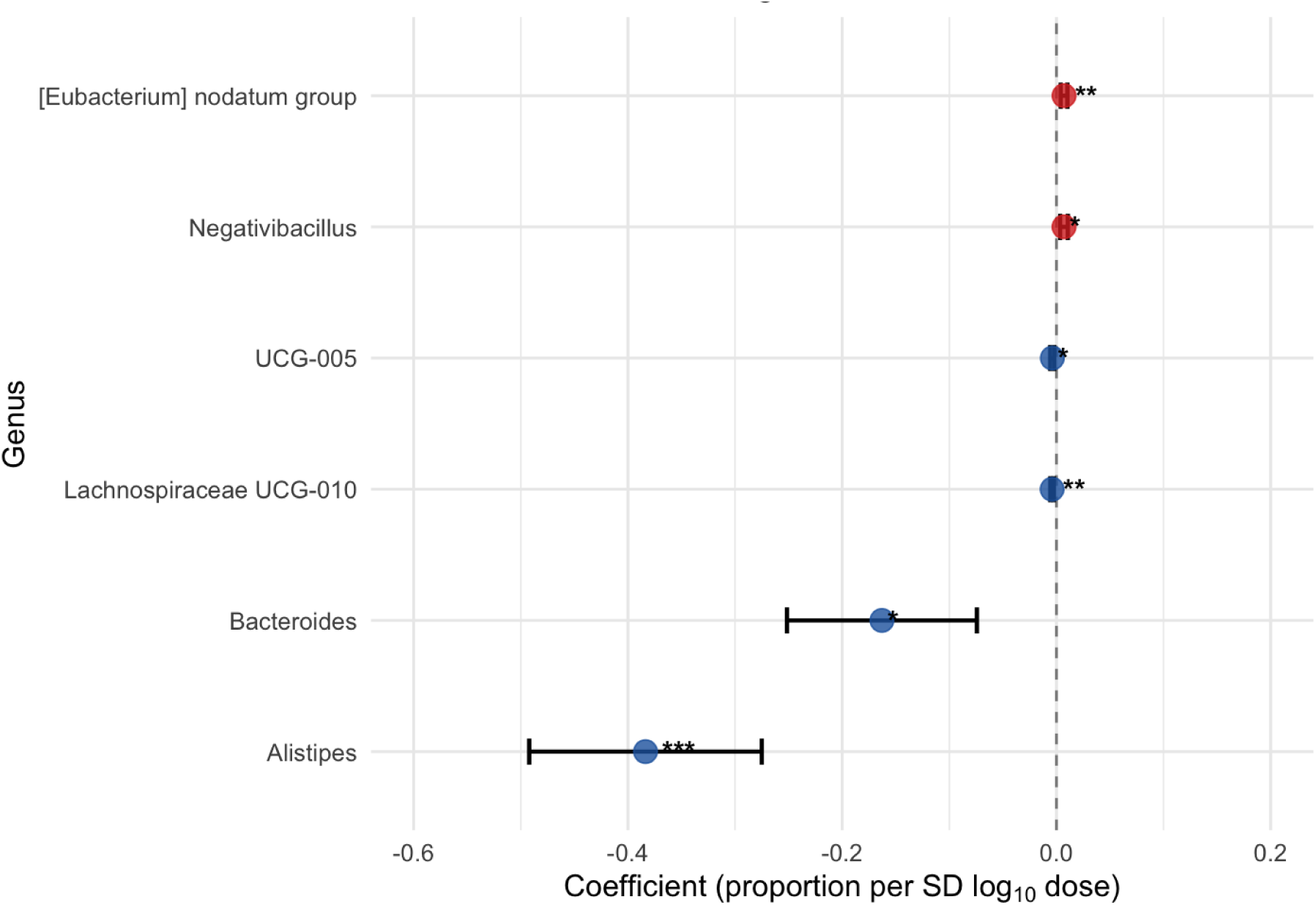
Significant dose-response effects on microbial genera. Linear dose-response analysis showing relationships between rapamycin dose and bacterial abundance changes across three studies (treated samples only; cumulative dose 42–990 mg/kg). Points represent linear model coefficients (proportion per SD log_10_ dose) with 95% confidence intervals. Red points indicate genera that increase with dose, while blue points show genera that decrease with dose. Seven genera showed significant dose-response associations (FDR < 0.05): *[Eubacterium] nodatum* group*\*\*,* and *Negativibacillus\** increase with dose, while UCG-005*\*,* Lachnospiraceae UCG-010*\*\*, Bacteroides\**, and *Alistipes*\**\**\* demonstrated depletion. The vertical dashed line represents no dose effect (coefficient = 0). Dose-response analysis was performed using MMUPHin batch-corrected data with linear models controlling for study effects. q < 0.001, ** q < 0.01, * q < 0.05.*

### Zero-inflated beta distributional meta-analysis

Having reported mean-level treatment effects after batch correction, we next examined distributional changes that act through both prevalence and non-zero abundance. We therefore fit zero-inflated beta GAMLSS models per study and combined the study coefficients in a random-effects meta-analysis. Following taxonomic harmonisation, 225 genera were retained for statistical analysis across all three studies. Zero-inflated beta regression models (BEZI family) were fitted with an overall success rate of 92.4%. Individual study analyses identified 35 genera with significant treatment effects at p < 0.05, distributed as 15 taxa in Bitto2016, 11 taxa in Han2021, and nine taxa in Yang2022. Applying within-study correction (FDR < 0.05), 16 taxa remained significant across studies.

For pooling, we required taxa to be present in ≥2 studies and to have stable estimates (SE ≤ 5), leaving 71 taxa eligible, with the metamicrobiomeR package applying additional internal filtering to yield 35 taxa in the final models. After FDR across the meta-analysed set, two genera showed significant treatments effects: *Bacteroides* demonstrated a pooled odds ratio (OR) of 1.59 (95% CI: 1.18–2.14, q = 0.036) with perfect cross-study consistency (I² = 0%). *Muribaculum* had an OR of 0.55 (95% CI: 0.38–0.80, q = 0.036) with moderate heterogeneity (I² = 68.4%). While *Ruminococcus* only showed trends (OR = 0.60, 95% CI: 0.41–0.89, q = 0.083) and displayed low heterogeneity (I² = 26.1%) (Fig.6; Table 2).

**Fig. 6.**
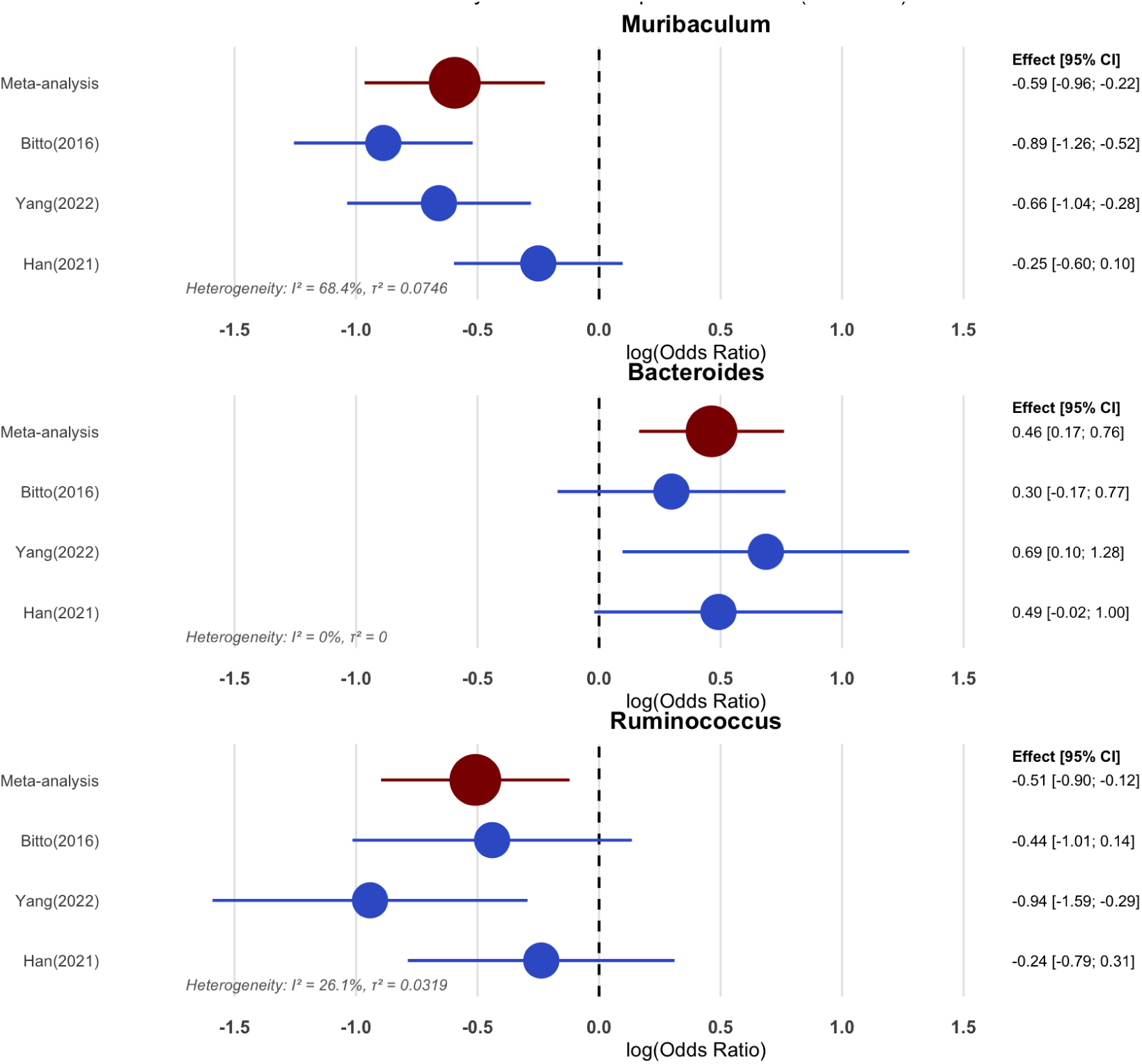
Forest plot with odds ratios for rapamycin treatment effects. Random-effects meta-analysis pooled results across three independent studies (Bitto et al., 2016; Yang et al., 2022; Han et al., 2021). The plot shows individual study estimates (blue) and the pooled effect (red) with 95% confidence intervals. Effect sizes are log-odds ratios from zero-inflated beta (BEZI) models; values > 0 indicate enrichment with rapamycin and values < 0 indicate depletion. After FDR correction, two genera were significant: *Muribaculum* −0.59 ± 0.19 (q = 0.042; k = 3) and *Bacteroides* +0.46 ± 0.15 (q = 0.042; k = 3). *Ruminococcus* showed a nominal decrease −0.51 ± 0.20 (q = 0.089; k = 3). Heterogeneity was low to moderate (*Bacteroides* I² = 0%; *Ruminococcus* I² = 26.1%; *Muribaculum* I² = 68.4%). Study weights were determined by inverse variance. Meta-analysis used metamicrobiomeR with GAMLSS-based modelling.

To examine dose sensitivity, Bitto2016 was subdivided by administration route, resulting in four study arms spanning a 23-fold dose range: Han2021 (42 mg/kg), Yang2022 (245 mg/kg), Bitto2016-IP (720 mg/kg) and Bitto2016-Diet (990 mg/kg). Meta-regression analysis of 35 taxa identified four genera with significant linear dose-response relationships (FDR < 0.05). *Lachnospira* exhibited the largest magnitude decline (slope = −0.00308 log OR per mg/kg, q = 4.92 × 10^-4^), corresponding to a 0.31% decrease in relative abundance per mg/kg. *A2* (family Lachnospiraceae) also decreased with dose (slope = −0.00058, q = 4.48 × 10^-3^), though the effect size was an order of magnitude smaller (0.058% per mg/kg). In contrast, *Faecalibaculum* increased with dose (slope = +0.0027, q = 3.37 × 10^-2^, +0.27% per mg/kg), and *Alistipes* showed a smaller positive trend (slope = +0.00077, q = 2.98 × 10^-2^). All significant associations had no detectable between-study heterogeneity (I² = 0%), indicating consistent dose-response patterns across the four arms. Slopes are expressed on the log-odds ratio scale from BEZI/GAMLSS estimates; multiplying the slope by the dose increment and exponentiating yields the corresponding odds-ratio change (Fig.7).

**Fig. 7:**
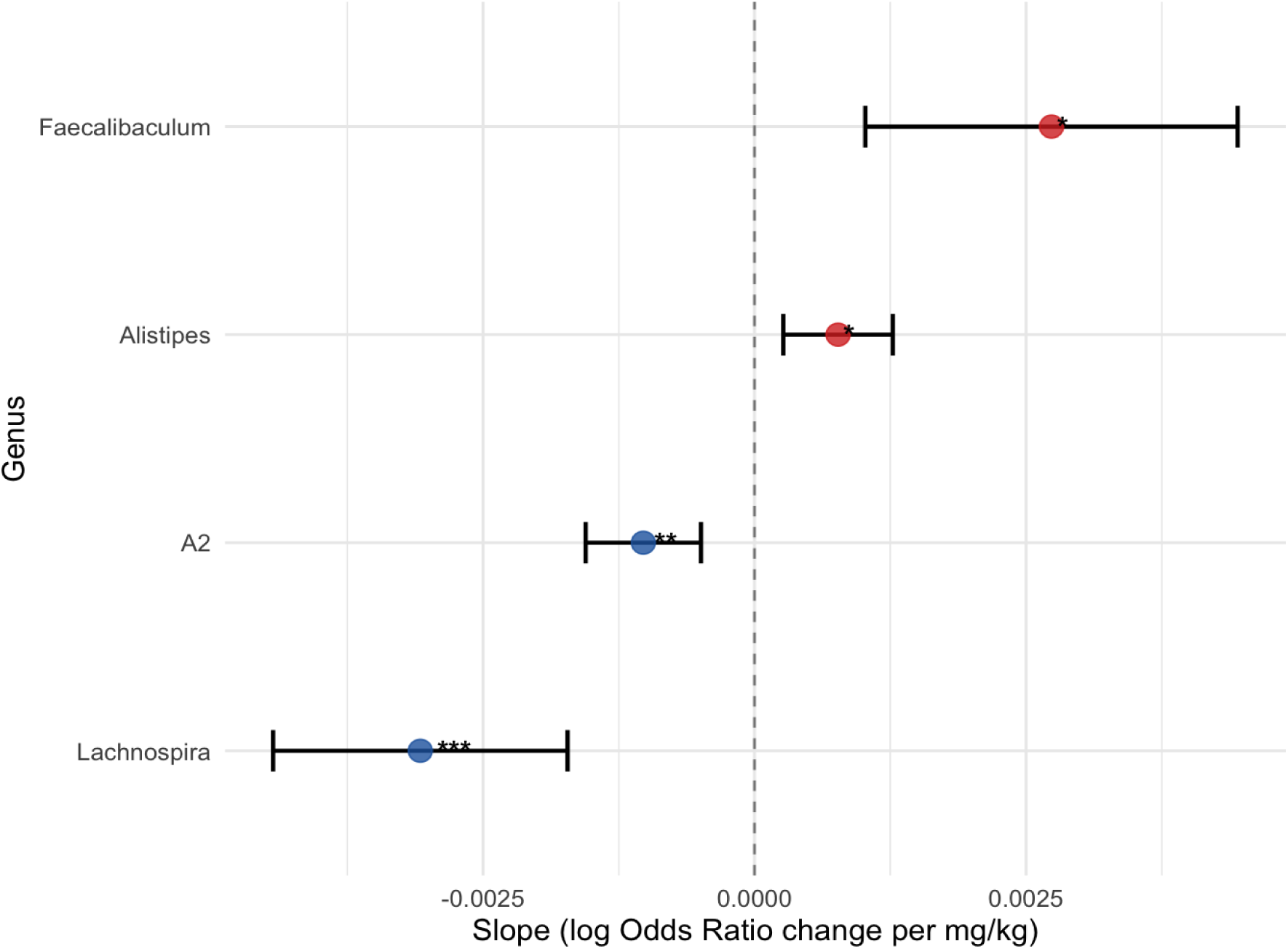
Odds ratio dose-response meta-regression rapamycin effects on gut bacterial genera. Meta-regression analysis showing dose-response relationships for rapamycin effects on gut bacterial genera across three studies. Points represent meta-regression slope coefficients (β; log odds ratio change per mg/kg dose) with 95% confidence intervals. Four genera showed dose-response associations (FDR < 0.05): *Lachnospira*\*\*\* demonstrates highly significant dose-related depletion, while Faecalibaculum* shows significant dose-related enrichment. *A2* (Lachnospiraceae)** and *Alistipes*\* showed weaker but significant dose-response effects. Meta-regression was performed using random-effects models in the metafor package with total dose (mg/kg) as a moderator variable. *** q < 0.001, ** q < 0.01, * q < 0.05.*

Excluding Bitto2016–Diet (990 mg/kg), we tested robustness across the IP-only range (42–720 mg/kg). In this subset, only *Alistipes* remained significant, with an effect size similar to the full range estimate; *Faecalibaculum* was not significant. *Lachnospira* was not evaluated because it was absent in Han2021, leaving fewer than three study arms with estimable effects (k = 2).

### Shared rapamycin signals across methods

Comparison of the GAMLSS and MMUPHin approaches revealed both convergent and complementary findings. Across both frameworks, *Bacteroides* was enriched and *Muribaculum* depleted with rapamycin (FDR < 0.05 in each), constituting cross-method evidence for these taxa.

Key methodological differences emerged in scope and sensitivity. GAMLSS analysed 71 genera with zero-inflated beta regression, identifying two significant taxa in meta-analysis (FDR < 0.05), while MMUPHin analysed 112 genera with batch-corrected linear models, identifying four significant taxa (FDR < 0.05). MMUPHin additionally detected ASF356 and *Ruminococcus* as significant, while GAMLSS showed *Ruminococcus* as marginally significant (q = 0.083).

For dose-response, GAMLSS modelled cumulative dose across the 42–990 mg/kg range and detected four genera with linear trends. At the same time, MMUPHin examined within-treatment dose gradients and identified six genera with significant associations. The two approaches converged on a core rapamycin-responsive set (*Bacteroides*, *Muribaculum*), while offering complementary strengths. MMUPHin controls inter-study batch structure and yields cross-study estimates, whereas GAMLSS explicitly models zero inflation and enables dose-response inference on the odds-ratio scale.

## Discussion

This study provides a cross-study synthesis of rapamycin-induced gut microbiome changes in mice, combining a standardised reanalysis of three independent 16S rRNA datasets with dual meta-analytic approaches and dose-response modelling. By applying consistent preprocessing and statistical frameworks, we identified reproducible rapamycin-associated microbial signatures, despite substantial methodological and biological differences between studies.

### Reanalysis confirms core findings while revealing methodological artefacts

Our reanalysis of Bitto et al. confirmed that rapamycin treatment alters community composition. Despite methodological ambiguity in their original approach (no specified distance metric or analytical details), by performing PERMANOVA using weighted UniFrac distances we obtained similar results in the injection cohort (our p = 0.020 vs. their p = 0.018), and results remained significant when pooling the IP and Diet cohorts (p = 0.012). However, our analysis in the feeding cohort showed only a trend (p = 0.096 vs. p = 0.015), suggesting greater variability or inadequate power with dietary rapamycin.

We observed higher relative abundances of segmented filamentous bacteria (SFB) Candidatus *Arthromitus* in rapamycin-treated groups across both delivery routes (Rapamycin–IP: 0.56% vs Control–IP: 0.25%; Rapamycin–Diet: 0.79% vs Control–Diet: 0.25%), consistent with original qPCR and histology findings. Although this directional trend matched the original study, differential abundance analysis using DESeq2 did not retain significance after FDR correction. Information from the Mouse Microbiome Database suggests that treated and control mice in this cohort had substantially elevated baseline SFB levels (10–40 times higher) compared to typical laboratory mice (Yang et al., 2019). This suggests that experimental conditions created an environmental predisposition to SFB colonisation, which rapamycin treatment may have amplified. The original study listed 16 taxa as significantly differentially abundant (p < 0.05), but we only found significant Muribaculaceae (formerly S24-7) enrichment after FDR-correction in the rapamycin IP group, potentially due to very low initial mean relative abundances (Lactococcus: 0.008%, Anaerostipes: 0.031%, Coprococcus: 0.007%). F/B ratios decreased significantly with rapamycin IP (0.67 vs 0.89; Wilcoxon p = 0.031) and trended downward with dietary rapamycin (0.83 vs 1.35; p = 0.130), consistent with the original violin-plot patterns and our observation of relative increases within Bacteroidetes.

In the Yang2022 dataset, which focused primarily on DSS-induced colitis in RhoB-deficient mice, potentially masking direct rapamycin effects, our reanalysis detected an alpha-diversity increase (Shannon index, p = 0.032) not reported in the original work, which was confirmed by community compositional shifts in unweighted UniFrac distances (PERMANOVA p = 0.009). We identified 31 significantly altered ASVs (FDR < 0.01), confirming the emergence of both Prevotellaceae and *Alloprevotella* lineages with rapamycin treatment. While several signals aligned with the original LEfSe analysis (*Helicobacter* and *Psychrobacter* increased; *Akkermansia* and Gastranaerophilales depleted), at the genus level, only *Helicobacter* remained significantly increased after FDR correction. The depletion of *Akkermansia* is particularly notable given its established role in promoting a healthy gut barrier, reducing inflammation, and protecting against metabolic disorders (Rodrigues et al., 2022). Notably, our analysis revealed ASV-level heterogeneity within *Muribaculum*, with some ASVs significantly enriched and others significantly depleted with rapamycin treatment, where LEfSe reported genus-level increase, underscoring the importance of ASV-level analysis over genus-level aggregation.

Han et al. investigated rapamycin effects and showed consistent beta-diversity shifts, which we confirmed by unweighted UniFrac analysis (PERMANOVA p = 0.007). We also found a decline in alpha-diversity(Shannon index p = 0.016) that the original study implied but did not explicitly test. Using LEfSe, the authors reported decreased abundances in beneficial (*Akkermansia*, Firmicutes-related groups) and increased abundances in potentially pathogenic taxa (*Helicobacter*, *Desulfovibrio*, *Sutterella*). However, we found that only *Helicobacter* showed significantly higher abundance after FDR correction (FDR < 0.01). Our analysis revealed ASV-level heterogeneity within genera, with bidirectional responses within *Alloprevotella*, Muribaculaceae-affiliated ASVs, and Lachnospiraceae NK4A136 group, with some ASVs increasing and others decreasing within the same genus.

The genus-level increase in *Helicobacter* in the Yang2022 and Han2021 datasets is worth noting. The potential for rapamycin to promote *Helicobacter* colonisation warrants careful consideration, particularly given the established role of the genus in gastric pathology (Sgamato et al., 2024). By contrast, the depletion of *Akkermansia* observed in both studies raises questions about potential trade-offs in the effects of rapamycin, as this beneficial mucin-degrader typically correlates with improved metabolic outcomes (Everard et al., 2013). However, effects are context- and strain-dependent, as they can also lead to mucus layer erosion and be vulnerable to inflammation (Luo et al., 2022). These opposing shifts, enrichment of a potentially pathogenic genus alongside depletion of a potentially beneficial one, highlight the complex ecological consequences of pharmacological mTOR inhibition on the gut microbiome.

### Cross-study core signals

Our standardised reanalysis identified several recurrent signals across datasets. Muribaculaceae enrichment emerged as the sole taxon that significantly increased in all three datasets. Additionally, consistent and statistically significant increases in *Helicobacter* abundance were observed in both Yang2022 and Han2021, supporting *Helicobacter* as a consistent rapamycin-associated responder under our analysis. The Lachnospiraceae NK4A136 group showed directionally similar abundance patterns in two studies.

Consistent declines in F/B ratios were observed across datasets, indicating a reproducible shift toward increased dominance of Bacteroidetes. Alpha diversity responses were varying across studies, where diversity significantly increased in Yang2022 (p = 0.032) and significantly decreased in Han2021 (p = 0.016), potentially linked to differences in experimental design such as dosage, administration route, or animal age.

By applying a uniform analytical framework, our reanalysis enhanced the detection and clarification of treatment effects that were present but not necessarily emphasised in the original analyses. The repeated emergence of Helicobacter across our reanalyses, and its enrichment reported previously (Bhat et al., 2017), underscores how standardisation surfaces shared signals that can be missed in single-study analyses. Additionally, applying systematic FDR correction demonstrated that, although original studies reported numerous taxa as significant, only a core subset withstands rigorous cross-study validation.

### Shared rapamycin signals across methods

We analysed the data using two complementary meta-analytic approaches: per-study zero-inflated beta modelling with random-effects pooling, and batch-corrected linear modelling. Both approaches identified increased *Bacteroides* and decreased *Muribaculum* as the most consistent rapamycin responses. *Bacteroides* enrichment is functionally significant due to its ability for SCFA production, including propionate and acetate, which fuel colonocyte metabolism and support gut barrier function (Horváth et al., 2022; Turnbaugh et al., 2006). Xu et al. (2020) found that rapamycin treatment increased *Bacteroides* abundance and SCFA levels, suggesting a mechanistic link.

*Bacteroides* enrichment further aligns with the consistent F/B ratio reductions we observed across studies. Wu et al. (2025) reported similar F/B shifts after 30 days of rapamycin treatment, alongside improved metabolic profiles. The *Bacteroides* enrichment across studies, independent of analytical methods, suggests rapamycin produces reproducible, potentially beneficial microbiome changes.

*Bacteroides* enrichment is particularly relevant to ageing biology. Ageing is associated with reduced abundance and diversity of protective anaerobes, including *Bacteroides* and *Bifidobacterium*, alongside shifts in dominant bacterial communities (Wilmanski et al., 2021; Zwielehner et al., 2009). Notably, centenarians exhibit distinct microbiome signatures, characterised by enriched *Bacteroidetes*, greater species evenness, and reduced pathobiont abundance compared to elderly individuals (Pang et al., 2023). Rapamycin-induced *Bacteroides* enrichment may contribute to healthier microbial ageing patterns, potentially supporting its broader effects on healthspan and lifespan.

While *Bacteroides* enrichment dominated the rapamycin response, we also observed consistent *Muribaculum* depletion across both meta-analytic frameworks. The decline of this taxon is notable given the role of *Muribaculum* as a SCFA producer (Ormerod et al., 2016; Lagkouvardos et al., 2019), albeit less prominent than *Bacteroides* in this capacity (Koh et al., 2016). In contrast to findings by Wu et al. (2025), who reported an increase in Muribaculum following rapamycin treatment, our meta-analysis identified consistent reductions across studies. These differences may reflect temporal dynamics, ASV-/strain-level heterogeneity within *Muribaculum,* or design differences across studies. Importantly, *Muribaculum* does not occur at the same abundance as *Bacteroides*, suggesting that its depletion may reflect functional reorganisation rather than a substantial loss of metabolic potential.

Across single-study analyses, pathway testing showed higher abundance of chorismate/aromatic-amino-acid biosynthesis (ALL-CHORISMATE-PWY), menaquinone (vitamin K_2_) biosynthesis (PWY-7373), and (p)ppGpp metabolism (PPGPPMET-PWY; stringent-response alarmone synthesis/turnover) under rapamycin. Because these functions are widespread across gut taxa, we interpret them as community-level shifts rather than effects attributable to a single genus. The (p)ppGpp signal is compatible with increased capacity for stress-adaptation; chorismate/AAA enrichment could support greater production of aromatic microbial metabolites (e.g., indoles); and menaquinone enrichment aligns with anaerobic electron-transport capacity. While some carbohydrate and amino-acid pathways also increased, inferring changes in mucosal oxygen from 16S-based pathway predictions alone cannot be made.

### Dose-response relationships

Dose-response analyses revealed that Lachnospiraceae taxa were the most consistently dose-sensitive. Lachnospiraceae UCG-010 (linear model) and *Lachnospira* and A2 (meta-regression) all showed dose-related depletion with zero heterogeneity (I² = 0%). Outside this family, results diverged depending on the method: the linear model indicated small *[Eubacterium] nodatum* group enrichment and *Alistipes* depletion, while meta-regression identified *Faecalibaculum* and *Alistipes* enrichment. Sensitivity analyses in meta regression, excluding the Bitto food arm as the highest dosage, showed that most associations were driven by the taxa mentioned above, without affecting *Allistipes*, suggesting that clinically relevant dose-response effects may be limited. Regardless, our results align well with the broader rapamycin microbiome literature. Effect sizes were small, suggesting that nominal dose explains slight variance compared with route, duration, and host context; we therefore treat dose-response findings as exploratory.

The effects of rapamycin on the microbiome are likely mediated by host intestinal epithelial changes. mTORC1 hyperactivation suppresses goblet and Paneth cell differentiation, whereas its inhibition restores these lineages (Zhou et al., 2015). Goblet cells produce mucus, a niche supporting mucin-degrading taxa such as *Bacteroides* and reduced *Akkermansia* (Png et al., 2010; Raimondi et al., 2021), while Paneth cells release antimicrobial peptides that preferentially target gram-positive taxa, including many Lachnospiraceae butyrate producers (Ryu et al., 2008; Sommer et al., 2014). This is compatible with the observed pattern of modest Lachnospiraceae declines alongside *Bacteroides* enrichment. Exposure kinetics (intraperitoneal bolus vs dietary delivery) likely exert more potent effects than mg/kg alone, given their distinct peak-trough tissue profiles (Trepanier et al., 1998).

The microbiome changes observed here occur within the established context of the longevity benefits of rapamycin (Miller et al., 2014; Bitto et al., 2016; Johnson & Kaeberlein, 2016; Johnson et al., 2015). Our data suggest a nuanced trade-off: depletion of butyrate-producing Lachnospiraceae could diminish anti-inflammatory capacity, yet enrichment of *Bacteroides*, a major SCFA and vitamin K producer, may preserve or even enhance certain host protective metabolic functions. These shifts occur alongside systemic mTOR inhibition, known to be associated with effects including reduced lipogenesis, enhanced mitochondrial biogenesis, and pyrimidine synthesis (Ben-Sahra et al., 2016; Saxton & Sabatini, 2017), and modulation of inflammatory pathways (Weichhart et al., 2015). Together, these observations suggest that rapamycin’s net outcome for host health will depend on the interplay between direct immune modulation, gut microbial metabolite production, and broader host metabolic reprogramming.

### Limitations

The primary limitation of our meta-analysis is the modest sample size (n=54 across three studies), which constrains the broader generalizability of our findings. Our cross-study meta-analytical approach necessarily integrates data from experiments with substantial methodological heterogeneity. The included studies differed in critical design parameters, including rapamycin dosing regimens, administration routes, treatment durations, and animal ages. Additionally, technical variations in DNA extraction protocols, primer selection (V4 versus V3–V4 regions), sequencing platforms, and taxonomic classification methods (BLAST against Greengenes versus RDP Classifier with SILVA) may have introduced systematic biases. These methodological differences can substantially affect taxonomic assignments (Escapa et al., 2020; Sierra, 2020) and reduce detected genera by up to 8% due to sequence truncation parameter variations (Abellan-Schneyder et al., 2021).

Accordingly, dose-sensitivity findings should be considered exploratory, and confirmatory experiments at intermediate doses and with harmonised exposure profiles will be important to characterise any non-linearities. Mechanistic interpretation of our findings is limited by multiple processes operating simultaneously, and our data alone cannot isolate specific pathways. Changes in goblet and Paneth cells under mTORC1 inhibition may shift mucus availability and antimicrobial-peptide pressure, but we lack direct measurements to detect these changes; therefore, these mechanisms remain speculative (Zhou et al., 2015; Ryu et al., 2008; Sommer et al., 2014).

Within these constraints, we identified recurrent rapamycin-associated microbiome signatures across the three heterogeneous studies, notably increased *Bacteroides* and decreased *Muribaculum*. However, given the differences in age, disease models, and dosing across studies, many secondary effects were context-dependent. With only three studies, we cannot capture the full variability in rapamycin-microbiome interactions, so additional studies with larger sample sizes and standardized protocols are needed to validate and expand upon these patterns.

## Conclusion

In summary, across three mouse datasets reanalysed with uniform methodology, rapamycin was associated with consistent but modest microbiome shifts characterised primarily by enrichment of *Bacteroides* and depletion of *Muribaculum*. While enrichment of *Bacteroides* aligns with patterns reported in healthy ageing cohorts and may contribute beneficial short-chain fatty acid production, concurrent depletion of other short-chain fatty acid producers and increases in *Helicobacter* highlight the ecological complexity of these changes. Alpha diversity responses varied by study context, and effects were influenced by dose, administration route, treatment duration and host factors.

Given the established longevity benefits of rapamycin alongside these complex microbiome effects, future clinical studies should incorporate microbiome monitoring and consider complementary interventions to maintain gut health during treatment. Adequately powered randomised studies with standardised protocols should include time course analyses, recovery phases after treatment cessation and arms that test prebiotic or probiotic co-interventions to enhance beneficial changes while mitigating potential risks. Functional readouts, including short-chain fatty acid profiles, metabolomics and barrier integrity measures, will be essential to link taxonomic shifts to health outcomes and to guide therapeutic strategies tenabling benefits of rapamycin in the gut microbiome and beyond.

## Supporting information

Supplementary Data: Additional Figures and Results

## Acknowledgements

We gratefully acknowledge funding support from Biovis

## Author Contributions

Conceptualization: IB; Data curation: AOM, IB; Resources: AOM, IB; Software: AOM, FM, IB; Methodology: AOM, FM, IB; Formal analysis: AOM; Investigation: AOM, IB; Validation: AOM, IB; Visualization: AOM, IB; Writing – original draft: AOM; Writing – review & editing: AOM, FM, PN, IZ, GF, IB; Funding acquisition: PN, IZ, GF; Supervision: GF, IB; Project administration: GF, IB.

## References

Abellan-Schneyder, I., Matchado, M. S., Reitmeier, S., Sommer, A., Sewald, Z., Baumbach, J., List, M., & Neuhaus, K. (2021). Primer, pipelines, parameters: Issues in 16S rRNA gene sequencing. mSphere, 6(1), e01202–20. 10.1128/mSphere.01202-20

Antoch, M. P., Wrobel, M., Gillard, B., Kuropatwinski, K. K., Toshkov, I., Gleiberman, A. S., Karasik, E., Moser, M. T., Foster, B. A., Andrianova, E. L., Chernova, O. V., & Gudkov, A. V. (2020). Superior cancer preventive efficacy of low versus high dose of mTOR inhibitor in a mouse model of prostate cancer. Oncotarget, 11(13), 1373–1387. 10.18632/oncotarget.27550

Baker, Darren J., Bennett G. Childs, Matej Durik, u. a. „Naturally Occurring p16Ink4a-Positive Cells Shorten Healthy Lifespan“. Nature 530, Nr. 7589 (2016): 184–89. 10.1038/nature16932.

Battesti, A., & Bouveret, E. (2009). Bacteria possessing two RelA/SpoT-like proteins have evolved a specific stringent response involving the acyl carrier protein-SpoT interaction. Journal of Bacteriology, 191(2), 616–624. 10.1128/JB.01195-08

Baker, D. J., Childs, B. G., Durik, M., Wijers, M. E., Sieben, C. J., Zhong, J., … van Deursen, J. M. (2016). Naturally occurring p16(Ink4a)-positive cells shorten healthy lifespan. Nature, 530(7589), 184–189. 10.1038/nature16932

Ben-Sahra, I., Hoxhaj, G., Ricoult, S. J. H., Asara, J. M., & Manning, B. D. (2016). mTORC1 induces purine synthesis through control of the mitochondrial tetrahydrofolate cycle. Science, 351(6274), 728–733. 10.1126/science.aad0489

Bhat, M., E. Pasini, J. Copeland, u. a. „Impact of Immunosuppression on the Metagenomic Composition of the Intestinal Microbiome: A Systems Biology Approach to Post-Transplant Diabetes“. Scientific Reports 7, Nr. 1 (2017): 10277. 10.1038/s41598-017-10471-2.

Bitto, A., Ito, T. K., Pineda, V. V., LeTexier, N. J., Huang, H. Z., Sutlief, E., Tung, H., Vizzini, N., Chen, B., Smith, K., Meza, D., Yajima, M., Beyer, R. P., … Kaeberlein, M. (2016). Transient rapamycin treatment can increase lifespan and healthspan in middle-aged mice. eLife, 5, e16351. 10.7554/eLife.16351

Bolyen, E., Rideout, J. R., Dillon, M. R., et al. (2019). Reproducible, interactive, scalable and extensible microbiome data science using QIIME 2. Nature Biotechnology, 37, 852–857. 10.1038/s41587-019-0209-9

Callahan, B. J., McMurdie, P. J., Rosen, M. J., Han, A. W., Johnson, A. J. A., & Holmes, S. P. (2016). DADA2: High-resolution sample inference from Illumina amplicon data. Nature Methods, 13(7), 581–583. 10.1038/nmeth.3869

Caspi, R., Billington, R., Keseler, I. M., Kothari, A., Krummenacker, M., Midford, P. E., Ong, W. K., Paley, S., Subhraveti, P., & Karp, P. D. (2020). The MetaCyc database of metabolic pathways and enzymes: A 2019 update. Nucleic Acids Research, 48(D1), D445–D453. 10.1093/nar/gkz862

Chenhuichen, Chenhui, Miriam Cabello-Olmo, Miguel Barajas, u. a. „Impact of Probiotics and Prebiotics in the Modulation of the Major Events of the Aging Process: A Systematic Review of Randomized Controlled Trials“. Experimental Gerontology 164 (Juli 2022): 111809. 10.1016/j.exger.2022.111809.

Cochran, William G. „The Combination of Estimates from Different Experiments“. Biometrics 10, Nr. 1 (1954): 101–29. 10.2307/3001666.

Conboy, I. M., Conboy, M. J., Wagers, A. J., Girma, E. R., Weissman, I. L., & Rando, T. A. (2005). Rejuvenation of aged progenitor cells by exposure to a young systemic environment. Nature, 433(7027), 760–764. 10.1038/nature03260

Costea, P. I., Zeller, G., Sunagawa, S., et al. (2017). Towards standards for human fecal sample processing in metagenomic studies. Nature Biotechnology, 35(11), 1069–1076. 10.1038/nbt.3960

Cruz-Bravo, R. K., Guevara-Gonzalez, R. G., Ramos-Gomez, M., Oomah, B. D., Wiersma, P., & Campos-Vega, R. (2014). The fermented non-digestible fraction of common bean (Phaseolus vulgaris L.) triggers cell cycle arrest and apoptosis in human colon adenocarcinoma cells. Genes & Nutrition, 9(1), 359. 10.1007/s12263-013-0359-1

DerSimonian, Rebecca, und Nan Laird. „Meta-analysis in clinical trials“. Controlled Clinical Trials 7, Nr. 3 (1986): 177–88. 10.1016/0197-2456(86)90046-2.

Davis, Cindy D. „The Gut Microbiome and Its Role in Obesity“. Nutrition Today 51, Nr. 4 (2016): 167–74. 10.1097/NT.0000000000000167.

Douglas, G. M., Maffei, V. J., Zaneveld, J. R., et al. (2020). PICRUSt2 for prediction of metagenome functions. Nature Biotechnology, 38(6), 685–688. 10.1038/s41587-020-0548-6

Escapa, I. F., Huang, Y., Chen, T., Lin, M., Kokaras, A., Dewhirst, F. E., & Lemon, K. P. (2020). Construction of habitat-specific training sets to achieve species-level assignment in 16S rRNA gene datasets. Microbiome, 8, 65. 10.1186/s40168-020-00841-w

Everard, Amandine, Clara Belzer, Lucie Geurts, u. a. „Cross-Talk between *Akkermansia Muciniphila* and Intestinal Epithelium Controls Diet-Induced Obesity“. Proceedings of the National Academy of Sciences 110, Nr. 22 (2013): 9066–71. 10.1073/pnas.1219451110.

Fan, W., Morinaga, H., Kim, J. J., Bae, E., Spann, N. J., Heinz, S., Glass, C. K., & Olefsky, J. M. (2010). FoxO1 regulates Tlr4 inflammatory pathway signalling in macrophages. The EMBO Journal, 29(24), 4223–4236. 10.1038/emboj.2010.274

Fok, Wilson C., Yidong Chen, Alex Bokov, u. a. „Mice fed rapamycin have an increase in lifespan associated with major changes in the liver transcriptome“. PloS one 9, Nr. 1 (2014): e83988. 10.1371/journal.pone.0083988.

Fujita, Y., Iki, M., Tamaki, J., Kouda, K., Yura, A., Kadowaki, E., … Kitamura, K. (2012). Association between vitamin K intake from fermented soybeans, natto, and bone mineral density in elderly Japanese men: The Fujiwara-kyo Osteoporosis Risk in Men (FORMEN) study. Osteoporosis International, 23(2), 705–714. 10.1007/s00198-011-1594-1

Gao, Yuan, und Tian Tian. „mTOR Signaling Pathway and Gut Microbiota in Various Disorders: Mechanisms and Potential Drugs in Pharmacotherapy“. International Journal of Molecular Sciences 24, Nr. 14 (2023): 11811. 10.3390/ijms241411811.

Guo, Jun, Xiuqing Huang, Lin Dou, u. a. „Aging and aging-related diseases: from molecular mechanisms to interventions and treatments“. Signal transduction and targeted therapy 7, Nr. 1 (2022): 391. 10.1038/s41392-022-01251-0.

Guo, Xue, Jing Xu, Chen Huang, u. a. „Rapamycin Extenuates Experimental Colitis by Modulating the Gut Microbiota“. Journal of Gastroenterology and Hepatology 38, Nr. 12 (2023): 2130–41. 10.1111/jgh.16381.

Han, W., Ma, Q., Liu, Y., et al. (2021). Sirolimus-induced metabolic disorders in mice are associated with intestinal microbiota dysbiosis, intestinal barrier dysfunction, and systemic inflammation. Transplantation, 105(5), 1017–1029. 10.1097/TP.0000000000003398

Harrison, P. W., Alako, B., Amid, C., et al. (2020). The European Nucleotide Archive in 2020. Nucleic Acids Research, 49(D1), D82–D85. 10.1093/nar/gkaa1028

Hipp, Mark S., Prasad Kasturi, und F. Ulrich Hartl. „The Proteostasis Network and Its Decline in Ageing“. Nature Reviews Molecular Cell Biology 20, Nr. 7 (2019): 421–35. 10.1038/s41580-019-0101-y.

Ho, N.T., Li, F., Wang, S. et al. metamicrobiomeR: an R package for analysis of microbiome relative abundance data using zero-inflated beta GAMLSS and meta-analysis across studies using random effects models. BMC Bioinformatics 20, 188 (2019). 10.1186/s12859-019-2744-2

Hopkins, M. J., Sharp, R., & Macfarlane, G. T. (2001). Age and disease related changes in intestinal bacterial populations assessed by cell culture, 16S rRNA abundance, and community cellular fatty acid profiles. Gut, 48(2), 198–205. 10.1136/gut.48.2.198

Horvath, T. D., Ihekweazu, F. D., Haidacher, S. J., Ruan, W., Engevik, K. A., Fultz, R., & Hoch, K. M. (2022). Bacteroides ovatus colonization influences the abundance of intestinal short chain fatty acids and neurotransmitters. iScience, 25(4), 104158. 10.1016/j.isci.2022.104158

Hurez, V., Dao, V., Liu, A., et al. (2015). Chronic mTOR inhibition in mice with rapamycin alters T, B, myeloid, and innate lymphoid cells and gut flora and prolongs life of immune-deficient mice. Aging Cell, 14(6), 945–956. 10.1111/acel.12380

Ivimey-Cook, Edward R., Zahida Sultanova, und Alexei A. Maklakov. „Rapamycin, Not Metformin, Mirrors Dietary Restriction-Driven Lifespan Extension in Vertebrates: A Meta-Analysis“. Aging Cell, 18. Juni 2025, e70131. 10.1111/acel.70131.

Jia, Kailiang, Di Chen, und Donald L. Riddle. „The TOR pathway interacts with the insulin signaling pathway to regulate C. elegans larval development, metabolism and life span“. *Development (Cambridge*, England) 131, Nr. 16 (2004): 3897–906. 10.1242/dev.01255.

Johnson, S. C., & Kaeberlein, M. (2016). Rapamycin in aging and disease: Maximizing efficacy while minimizing side effects. Oncotarget, 7(29), 44876–44878. 10.18632/oncotarget.10381

Johnson, S. C., Yanos, M. E., Bitto, A., Castanza, A., Gagnidze, A., Gonzalez, B., Gupta, K., Hui, J., Jarvie, C., Johnson, B. M., Letexier, N., McCanta, L., Sangesland, M., … Kaeberlein, M. (2015). Dose-dependent effects of mTOR inhibition on weight and mitochondrial disease in mice. Frontiers in Genetics, 6, 247. 10.3389/fgene.2015.00247

Jung, M. J., Lee, J., Shin, N. R., et al. (2016). Chronic repression of mTOR Complex 2 induces changes in the gut microbiota of diet-induced obese mice. Scientific Reports, 6, 30887. 10.1038/srep30887

Kapahi, Pankaj, Brian M. Zid, Tony Harper, Daniel Koslover, Viveca Sapin, und Seymour Benzer. „Regulation of lifespan in Drosophila by modulation of genes in the TOR signaling pathway“. Current biology : CB 14, Nr. 10 (2004): 885–90. 10.1016/j.cub.2004.03.059.

Kennedy, B. K., & Lamming, D. W. (2016). The mechanistic target of rapamycin: The grand conductor of metabolism and aging. Cell Metabolism, 23(6), 990–1003. 10.1016/j.cmet.2016.05.009

Koh, A., De Vadder, F., Kovatcheva-Datchary, P., & Bäckhed, F. (2016). From dietary fiber to host physiology: Short-chain fatty acids as key bacterial metabolites. Cell, 165(6), 1332–1345. 10.1016/j.cell.2016.05.041

Lagkouvardos, I., Lesker, T. R., Hitch, T. C. A., Gálvez, E. J. C., Smit, N., Neuhaus, K., Wang, J., Baines, J. F., Abt, B., Stecher, B., Overmann, J., Strowig, T., & Clavel, T. (2019). Sequence and cultivation study of Muribaculaceae reveals novel species, host preference, and functional potential of this yet undescribed family. Microbiome, 7(1), 1–15. 10.1186/s40168-019-0637-2

Lamming, D. W., Ye, L., Katajisto, P., Goncalves, M. D., Saitoh, M., Stevens, D. M., … Sabatini, D. M. (2012). Rapamycin-induced insulin resistance is mediated by mTORC2 loss and uncoupled from longevity. Science, 335(6076), 1638–1643. 10.1126/science.1215135

Lau, Agnes Wei Yin, Loh Teng-Hern Tan, Nurul-Syakima Ab Mutalib, Sunny Hei Wong, Vengadesh Letchumanan, und Learn-Han Lee. „The chemistry of gut microbiome in health and diseases“. Progress In Microbes & Molecular Biology 4, Nr. 1 (2021). 10.36877/pmmb.a0000175.

Lee, Deborah J W, Ajla Hodzic Kuerec, und Andrea B Maier. „Targeting Ageing with Rapamycin and Its Derivatives in Humans: A Systematic Review“. The Lancet Healthy Longevity 5, Nr. 2 (2024): e152–62. 10.1016/S2666-7568(23)00258-1.

Leinonen, R., Sugawara, H., & Shumway, M. (2011). The Sequence Read Archive. Nucleic Acids Research, 39(suppl_1), D19–D21. 10.1093/nar/gkq1019

Leo Lahti [Aut, Cre]. microbiome. Bioconductor, released 2017. 10.18129/B9.BIOC.MICROBIOME.

Lim, Shaun H. Y., Malene Hansen, und Caroline Kumsta. „Molecular Mechanisms of Autophagy Decline during Aging“. Cells 13, Nr. 16 (2024): 1364. 10.3390/cells13161364.

Li, J., Kim, S. G., & Blenis, J. (2014). Rapamycin: One drug, many effects. Cell Metabolism, 19(3), 373–379. 10.1016/j.cmet.2014.01.001

Lin, A. L., Aware, C., Neher, C., Hamdi, M., Ericsson, A., Khegai, O., Patrie, J., Kurt, M., Govindarajan, M., Woods, C., Ivanich, K., Beversdorf, D., Cheng, J., Balchandani, P., Gonzales, M., & Altes, T. (2025). Rapamycin enhances neurovascular, peripheral metabolic, and immune function in cognitively normal, middle-aged APOE4 carriers: Genotype-dependent effects compared to non-carriers. Research Square. 10.21203/rs.3.rs-6214340/v1

López-Otín, Carlos, Maria A. Blasco, Linda Partridge, Manuel Serrano, und Guido Kroemer. „Hallmarks of aging: An expanding universe“. Cell 186, Nr. 2 (2023): 243–78. 10.1016/j.cell.2022.11.001.

Love, M. I., Huber, W., & Anders, S. (2014). Moderated estimation of fold change and dispersion for RNA-seq data with DESeq2. Genome Biology, 15(12), 550. 10.1186/s13059-014-0550-8

Luo, Yuheng, Cong Lan, Hua Li, u. a. „Rational Consideration of Akkermansia Muciniphila Targeting Intestinal Health: Advantages and Challenges“. Npj Biofilms and Microbiomes 8, Nr. 1 (2022): 81. 10.1038/s41522-022-00338-4.

Manes, Annalaura, Tiziana Di Renzo, Loreta Dodani, u. a. „Pharmacomicrobiomics of Classical Immunosuppressant Drugs: A Systematic Review“. Biomedicines 11, Nr. 9 (2023): 2562. 10.3390/biomedicines11092562.

Mannick, Joan B, Grace Teo, Patti Bernardo, u. a. „Targeting the Biology of Ageing with mTOR Inhibitors to Improve Immune Function in Older Adults: Phase 2b and Phase 3 Randomised Trials“. The Lancet Healthy Longevity 2, Nr. 5 (2021): e250–62. 10.1016/S2666-7568(21)00062-3.

McHugh, Domhnall, und Jesús Gil. „Senescence and Aging: Causes, Consequences, and Therapeutic Avenues“. Journal of Cell Biology 217, Nr. 1 (2018): 65–77. 10.1083/jcb.201708092.

McMurdie, P. J., & Holmes, S. (2013). phyloseq: An R package for reproducible interactive analysis and graphics of microbiome census data. PLOS ONE, 8(4), e61217. 10.1371/journal.pone.0061217

Meiners, Franziska, Bernd Kreikemeyer, Patrick Newels, u. a. „Strawberry Dietary Intervention Influences Diversity and Increases Abundances of SCFA-Producing Bacteria in Healthy Elderly People“. Microbiology Spectrum 13, Nr. 2 (2025): e01913–24. 10.1128/spectrum.01913-24.

Meiners, Franziska, Asiri Ortega-Matienzo, Georg Fuellen, und Israel Barrantes. „Gut microbiome-mediated health effects of fiber and polyphenol-rich dietary interventions“. Frontiers in Nutrition 12 (August 2025): 1647740. 10.3389/fnut.2025.1647740.

Miller, R. A., Harrison, D. E., Astle, C. M., Fernandez, E., Flurkey, K., Han, M., Javors, M. A., Li, X., Nadon, N. L., Nelson, J. F., Pletcher, S., Salmon, A. B., Sharp, Z. D., … Strong, R. (2014). Rapamycin-mediated lifespan increase in mice is dose and sex dependent and metabolically distinct from dietary restriction. Aging Cell, 13(3), 468–477. 10.1111/acel.12194

Moqri, M., Herzog, C., Poganik, J. R., Biomarkers of Aging Consortium, Justice, J., Belsky, D. W., Higgins-Chen, A., Moskalev, A., Fuellen, G., Cohen, A. A., Bautmans, I., Widschwendter, M., Ding, J., Fleming, A., Mannick, J., Han, J.-D. J., Zhavoronkov, A., Barzilai, N., Kaeberlein, M., … Gladyshev, V. N. (2023). Biomarkers of aging for the identification and evaluation of longevity interventions. Cell, 186(18), 3758–3775. 10.1016/j.cell.2023.07.003

Neff, F., Flores-Dominguez, D., Ryan, D. P., Horsch, M., Schröder, S., Adler, T., … Ehninger, G. (2013). Rapamycin extends murine lifespan but has limited effects on aging. Journal of Clinical Investigation, 123(8), 3272–3291. 10.1172/JCI67674

Niccoli, Teresa, und Linda Partridge. „Ageing as a risk factor for disease“. Current biology : CB 22, Nr. 17 (2012): 17. 10.1016/j.cub.2012.07.024.

Oksanen, J., Blanchet, F. G., Friendly, M., et al. (2018). vegan: Community ecology package (Version 2.5-2) [R package].

Noureldein, Mohamed H., und Assaad A. Eid. „Gut microbiota and mTOR signaling: Insight on a new pathophysiological interaction“. Microbial pathogenesis 118 (2018): 98–104. 10.1016/j.micpath.2018.03.021.

Ormerod, K. L., Wood, D. L., Lachner, N., Gellatly, S. L., Daly, J. N., Parsons, J. D., Dal’Molin, C. G. O., Palfreyman, R. W., Nielsen, L. K., Cooper, M. A., Morrison, M., Hansbro, P. M., & Hugenholtz, P. (2016). Genomic characterization of the uncultured Bacteroidales family S24-7 inhabiting the guts of homeothermic animals. Microbiome, 4(1), 36. 10.1186/s40168-016-0181-2

Pang, S., Chen, X., Lu, Z., Meng, L., Huang, Y., Yu, X., et al. (2023). Longevity of centenarians is reflected by the gut microbiome with youth-associated signatures. Nature Aging, 3, 436–449. 10.1038/s43587-023-00455-5

Papadopoli D, Boulay K, Kazak L et al. mTOR as a central regulator of lifespan and aging [version 1; peer review: 3 approved]. F1000Research 2019, 8(F1000 Faculty Rev):998 (10.12688/f1000research.17196.1)

Partridge, L., Fuentealba, M., & Kennedy, B. K. (2020). The quest to slow ageing through drug discovery. Nature Reviews Drug Discovery, 19(8), 513–532. 10.1038/s41573-020-0067-7

Paulson JN, Stine OC, Bravo HC, Pop M (2013). “Differential abundance analysis for microbial marker-gene surveys.” Nat Meth, advance online publication. doi:10.1038/nmeth.2658

Powers, R. Wilson, Matt Kaeberlein, Seth D. Caldwell, Brian K. Kennedy, und Stanley Fields. „Extension of chronological life span in yeast by decreased TOR pathway signaling“. Genes & development 20, Nr. 2 (2006): 174–84. 10.1101/gad.1381406.

Price, M. N., Dehal, P. S., & Arkin, A. P. (2010). FastTree 2 – Approximately maximum-likelihood trees for large alignments. PLOS ONE, 5(3), e9490. 10.1371/journal.pone.0009490

Quast, C., Pruesse, E., Yilmaz, P., et al. (2013). The SILVA ribosomal RNA gene database project: Improved data processing and web-based tools. Nucleic Acids Research, 41(D1), D590–D596. 10.1093/nar/gks1219

Ravikrishnan, Aarthi, Indrik Wijaya, Eileen Png, u. a. „Gut Metagenomes of Asian Octogenarians Reveal Metabolic Potential Expansion and Distinct Microbial Species Associated with Aging Phenotypes“. Nature Communications 15, Nr. 1 (2024): 7751. 10.1038/s41467-024-52097-9.

Rios-Covián, D., Arboleya, S., Hernandez-Barranco, A. M., Alvarez-Buylla, J. R., Ruas-Madiedo, P., & Gueimonde, M. (2013). Interactions between Bifidobacterium and Bacteroides species in cofermentations are affected by carbon sources, including exopolysaccharides produced by bifidobacteria. Applied and Environmental Microbiology, 79(23), 7518–7524. 10.1128/AEM.02545-13

Rios-Covián, D., Sánchez, B., Salazar, N., Martinez, N., Redruello, B., Gueimonde, M., & de los Reyes-Gavilán, C. G. (2015). Different metabolic features of Bacteroides fragilis growing in the presence of glucose and exopolysaccharides of bifidobacteria. Frontiers in Microbiology, 6, 825. 10.3389/fmicb.2015.00825

Rodrigues, V. F.; Elias-Oliveira, J.; Pereira, Í. S.; Pereira, J. A.; Barbosa, S. C.; Machado, M. S. G.; Carlos, D. Akkermansia muciniphila and Gut Immune System: A Good Friendship That Attenuates Inflammatory Bowel Disease, Obesity, and Diabetes. Frontiers in immunology 2022, 13, 934695. DOI: 10.3389/fimmu.2022.934695.

Ryu, J.-H., Kim, S.-H., Lee, H.-Y., Bai, J. Y., Nam, Y.-D., Bae, J.-W., Lee, D. G., Shin, S. C., Ha, E.-M., & Lee, W.-J. (2008). Innate immune homeostasis by the homeobox gene caudal and commensal-gut mutualism in Drosophila. Science, 319(5864), 777–782. 10.1126/science.1149357

Saxton, R. A., & Sabatini, D. M. (2017). mTOR signaling in growth, metabolism, and disease. Cell, 168(6), 960–976. 10.1016/j.cell.2017.02.004

Sgamato C, Rocco A, Compare D, Priadko K, Romano M, Nardone G. Exploring the Link between Helicobacter pylori, Gastric Microbiota and Gastric Cancer. Antibiotics (Basel). 2024 May 24;13(6):484. doi: 10.3390/antibiotics13060484. PMID: 38927151; PMCID: PMC11201017.

Shimizu, J., Kubota, T., Takada, E., Takai, K., Fujiwara, N., & Arimitsu, N. (2018). Propionate-producing bacteria in the intestine may associate with skewed responses of IL10-producing regulatory T cells in patients with relapsing polychondritis. PLOS ONE, 13(9), e0203657. 10.1371/journal.pone.0203657

Sierra, M. A., Li, Q., Pushalkar, S., Paul, B., Sandoval, T. A., Kamer, A. R., Corby, P., Guo, Y., Ruff, R. R., Alekseyenko, A. V., Li, X., & Saxena, D. (2020). The influences of bioinformatics tools and reference databases in analyzing the human oral microbial community. Genes, 11(8), 878. 10.3390/genes11080878

Sinha, Anurag Kumar, Martin Frederik Laursen, und Tine Rask Licht. „Regulation of Microbial Gene Expression: The Key to Understanding Our Gut Microbiome“. Trends in Microbiology, August 2024, S0966842X24001756. 10.1016/j.tim.2024.07.005.

Smith, P., Willemsen, D., Popkes, M., Metge, F., Gandiwa, E., Reichard, M., & Valenzano, D. R. (2022). Regulation of life span by the gut microbiota in the short-lived African turquoise killifish. eLife, 6, e27014. 10.7554/eLife.27014

Sommer, Felix, Nina Adam, Malin E. V. Johansson, Lijun Xia, Gunnar C. Hansson, und Fredrik Bäckhed. „Altered Mucus Glycosylation in Core 1 O-Glycan-Deficient Mice Affects Microbiota Composition and Intestinal Architecture“. PLoS ONE 9, Nr. 1 (2014): e85254. 10.1371/journal.pone.0085254.

Sprott, Richard L. „Biomarkers of aging and disease: introduction and definitions“.Experimental gerontology 45, Nr. 1 (2010): 1. 10.1016/j.exger.2009.07.008.

Sun, Jiahui, Xiaoxuan Wang, Junhong Xiao, u. a. „Autophagy Mediates the Impact of Porphyromonas Gingivalis on Short-Chain Fatty Acids Metabolism in Periodontitis-Induced Gut Dysbiosis“. Scientific Reports 14, Nr. 1 (2024): 26291. 10.1038/s41598-024-77909-2.

Stasinopoulos, M. (2012). gamlss: Generalized Additive Models for Location, Scale and Shape (R package version 5.4–22). https://cran.r-project.org/web/packages/gamlss

Trepanier, D. J., Gallant, H., Legatt, D. F., & Yatscoff, R. W. (1998). Rapamycin: Distribution, pharmacokinetics and therapeutic range investigations: An update. Clinical Biochemistry, 31(5), 345–351. 10.1016/S0009-9120(98)00048-4

Turnbaugh, P. J., Ley, R. E., Mahowald, M. A., Magrini, V., Mardis, E. R., & Gordon, J. I. (2006). An obesity-associated gut microbiome with increased capacity for energy harvest. Nature, 444(7122), 1027–1031.

Vinolo, M. A. R., Rodrigues, H. G., Nachbar, R. T., & Curi, R. (2011). Regulation of inflammation by short chain fatty acids. Nutrients, 3(10), 858–876. 10.3390/nu3100858

Walther, B., Karl, J. P., Booth, S. L., & Boyaval, P. (2013). Menaquinones, bacteria, and the food supply: The relevance of dairy and fermented food products to vitamin K requirements. Advances in Nutrition, 4(4), 463–473. 10.3945/an.113.003855

Weichhart, T., Hengstschlager, M., & Linke, M. (2015). Regulation of innate immune cell function by mTOR. Nature Reviews Immunology, 15(10), 599–614. 10.1038/nri3901

Wilmanski, Tomasz, Christian Diener, Noa Rappaport, u. a. „Gut microbiome pattern reflects healthy ageing and predicts survival in humans“. Nature metabolism 3, Nr. 2 (2021): 274–86. 10.1038/s42255-021-00348-0).

Wu, L., Kensiski, A., Gavzy, S. J., Lwin, H. W., Song, Y., France, M. T., Lakhan, R., Kong, D., Li, L., Saxena, V., Piao, W., Shirkey, M. W., Mas, V. R., Ma, B., & Bromberg, J. S. (2025). Rapamycin immunomodulation utilizes time-dependent alterations of lymph node architecture, leukocyte trafficking, and gut microbiome. JCI Insight, 10(8), e186505. 10.1172/jci.insight.186505

Xu, L., Zhang, C., He, D., Jiang, N., Bai, Y., & Xin, Y. (2020). Rapamycin and MCC950 modified gut microbiota in experimental autoimmune encephalomyelitis mouse by brain gut axis. Life Sciences, 253, 117747. 10.1016/j.lfs.2020.117747

Yan, Haoteng, Jie Ren, und Guang-Hui Liu. „Fecal Microbiota Transplantation: A New Strategy to Delay Aging“. hLife 1, Nr. 1 (2023): 8–11. 10.1016/j.hlife.2023.06.002.

Yang, J., Park, J., Park, S., Baek, I., & Chun, J. (2019). Introducing Murine Microbiome Database (MMDB): A Curated Database with Taxonomic Profiling of the Healthy Mouse Gastrointestinal Microbiome. Microorganisms, 7(11), 480. 10.3390/microorganisms7110480

Yang, J., Pei, G., Sun, X. et al. RhoB affects colitis through modulating cell signaling and intestinal microbiome. Microbiome 10, 149 (2022). 10.1186/s40168-022-01347-3

Zhang, Guolin, Yuqing Lu, Zhen Wang, u. a. „Causal relationship between gut microbiota and ageing: A multi-omics Mendelian randomization study“. Archives of Gerontology and Geriatrics 131 (April 2025): 105765. 10.1016/j.archger.2025.105765.

Zhang, Yong, Qiangchuan Hou, Chen Ma, u. a. „Lactobacillus casei protects dextran sodium sulfate-or rapamycin-induced colonic inflammation in the mouse“. European journal of nutrition 59, Nr. 4 (2020): 1443–51. 10.1007/s00394-019-02001-9.

Zhao, Yang, Matthew Simon, Andrei Seluanov, und Vera Gorbunova. „DNA Damage and Repair in Age-Related Inflammation“. Nature Reviews Immunology 23, Nr. 2 (2023): 75–89. 10.1038/s41577-022-00751-y.

Zhou Y, Rychahou P, Wang Q, Weiss HL, Evers BM. TSC2/mTORC1 signaling controls Paneth and goblet cell differentiation in the intestinal epithelium. Cell Death Dis. 2015 Feb 5;6(2):e1631. doi: 10.1038/cddis.2014.588.

Zwielehner, Jutta, Kathrin Liszt, Michael Handschur, Cornelia Lassl, Alexander Lapin, und Alexander G. Haslberger. „Combined PCR-DGGE Fingerprinting and Quantitative-PCR Indicates Shifts in Fecal Population Sizes and Diversity of Bacteroides, Bifidobacteria and Clostridium Cluster IV in Institutionalized Elderly“. Experimental Gerontology 44, Nr. 6–7 (2009): 440–46. 10.1016/j.exger.2009.04.002.

