## Supplementary Data: Additional Figures and Results for "Meta-analysis reveals reproducible rapamycin-induced shifts in the mouse gut microbiome"

### Supplements First Draft

#### Taxonomic Composition Analysis

##### Phylum-Level Composition

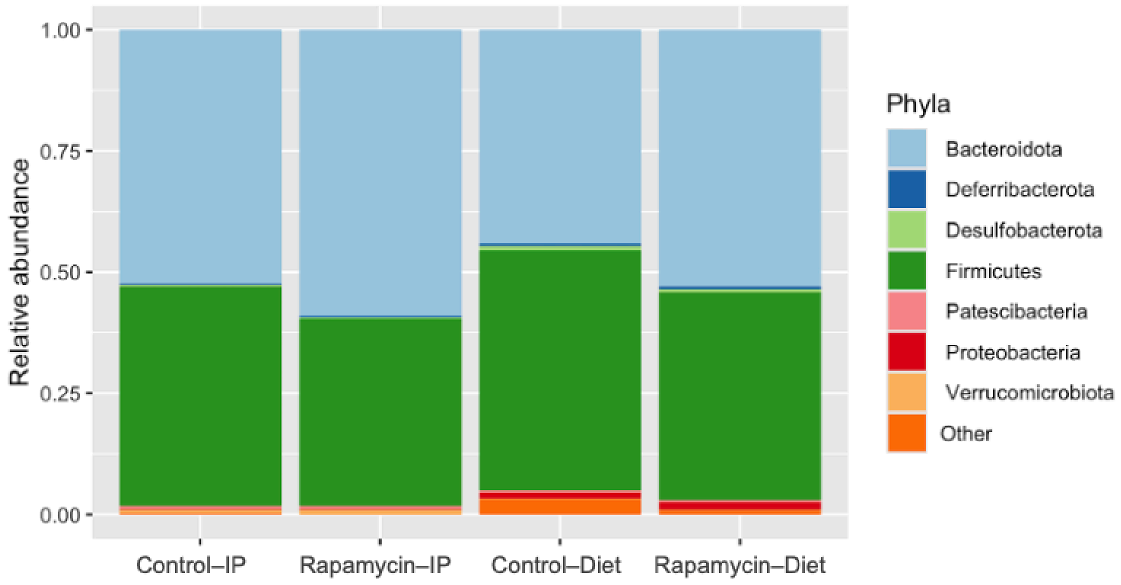


**Fig. S1 Relative abundance of bacterial phyla** across rapamycin treatment groups in the Bitto2016 dataset. Stacked bar chart showing the proportional representation of major bacterial phyla in control and rapamycin-treated groups across intraperitoneal (IP) and dietary administration routes. Firmicutes and Bacteroidetes represent the dominant phyla across all treatment conditions, comprising approximately 80-90% of gut microbial communities. No statistically significant differences in phylum-level composition were observed between rapamycin-treated and control groups (Kruskal-Wallis test, p = 0.7168). Each bar represents 100% relative abundance with individual phyla contributions shown by colored segments according to the legend

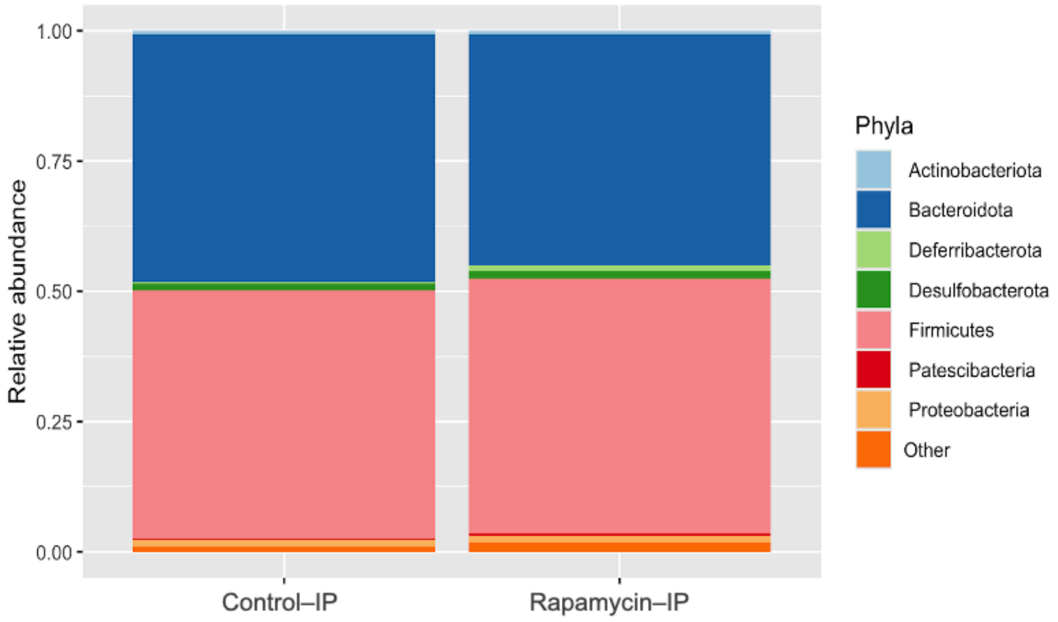


**Fig. S2. Relative abundance of bacterial phyla** across rapamycin treatment groups in the Yang2022 dataset. Stacked bar chart showing the proportional representation of major bacterial phyla in control and rapamycin-treated groups with intraperitoneal (IP) administration. Firmicutes and Bacteroidetes represent the dominant phyla across both treatment conditions, comprising approximately 80-90% of gut microbial communities. No statistically significant differences in phylum-level composition were observed between rapamycin-treated and control groups (Kruskal-Wallis test, $p = 0.4622$). Each bar represents 100% relative abundance with individual phyla contributions shown by colored segments according to the legend

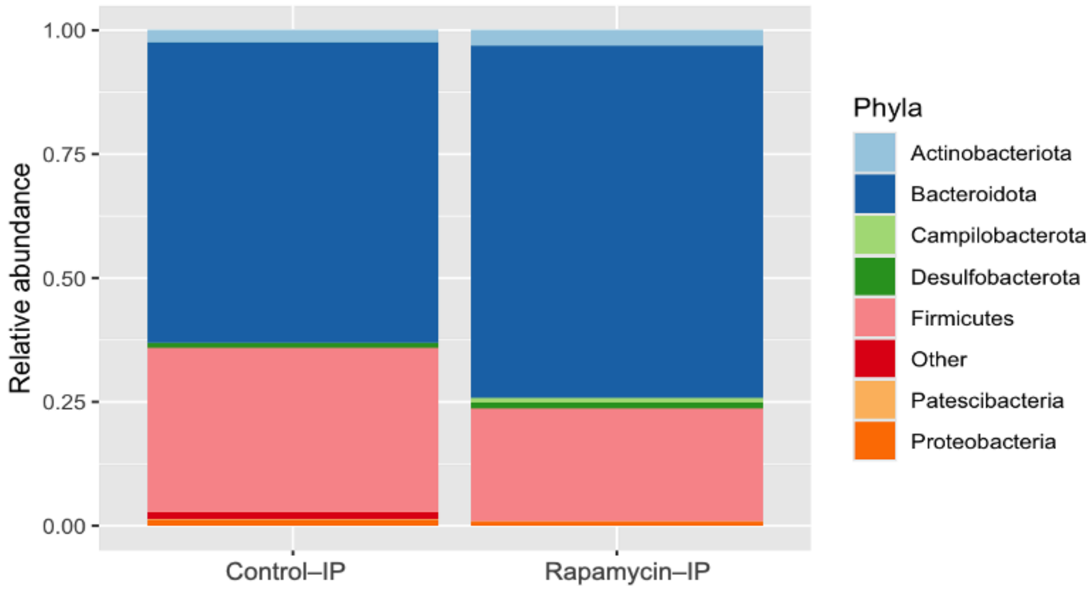


**Fig. S3 Relative abundance of bacterial phyla** across rapamycin treatment groups in the Han et al. dataset. Stacked bar chart showing the proportional representation of major bacterial phyla in control and rapamycin-treated groups with intraperitoneal (IP) administration. Firmicutes and Bacteroidetes represent the dominant phyla across both treatment conditions, comprising approximately 80-90% of gut microbial communities. No statistically significant differences in phylum-level composition were observed between rapamycin-treated and control groups

(Kruskal-Wallis test, p = 0.8847). Each bar represents 100% relative abundance with individual phyla contributions shown by colored segments according to the legend

#### Genus-Level Composition

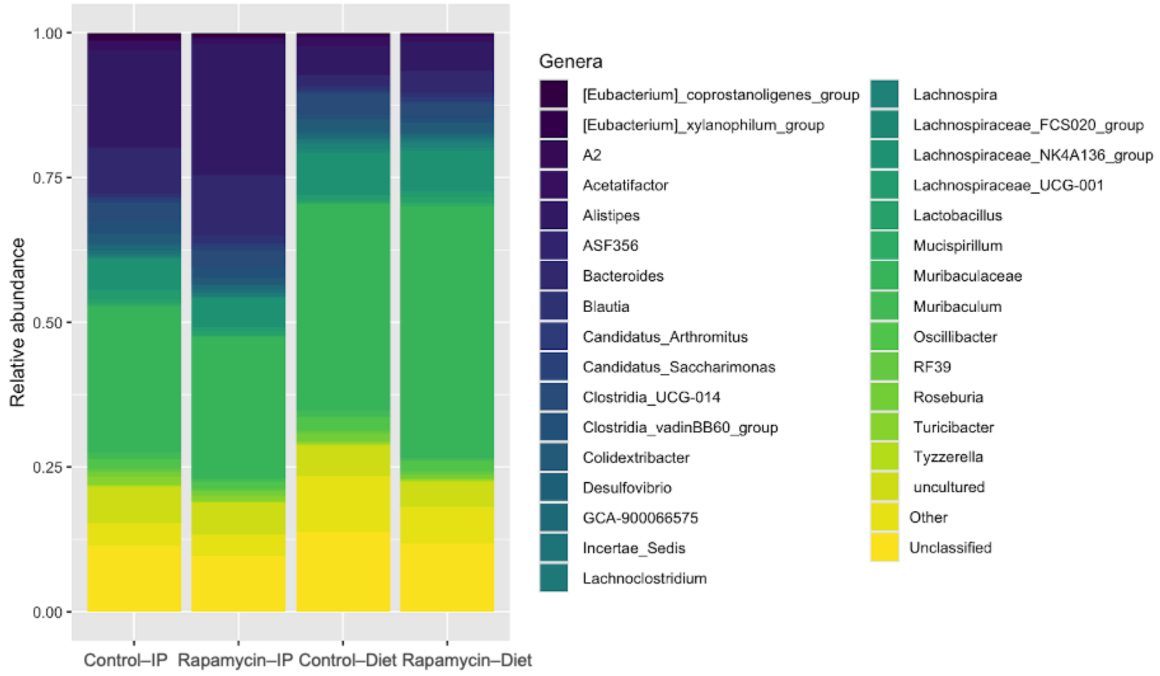


**Fig. S4** Genus composition
Bitto2016 (IP and Diet); stacked bars show mean relative abundance per group (Control–IP, Rapamycin–IP, Control–Diet, Rapamycin–Diet); colours denote genera; inclusion thresholds for this panel were detection 1.25% (relative abundance) and prevalence ≥50%; "Other" pools low-abundance genera and "Unclassified" denotes reads not assigned at genus level; genus-level composition did not differ across groups (Kruskal–Wallis p = 0.9728)

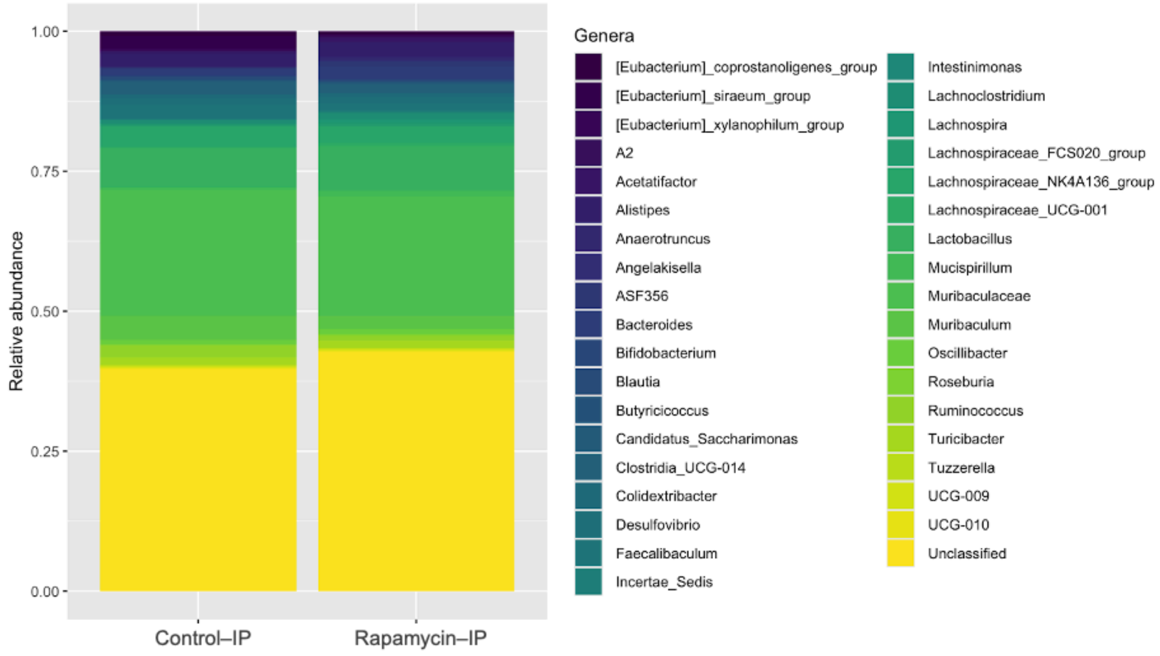


**Fig. S5 Genus composition**

Yang2022 (IP); stacked bars show mean relative abundance per group (Control–IP, Rapamycin–IP); colours denote genera; inclusion thresholds were detection 1.25% (relative abundance) and prevalence ≥50%; “Other” pools low-abundance genera and “Unclassified” denotes reads not assigned at genus level; genus-level composition did not differ between groups (Kruskal–Wallis p = 0.5061)

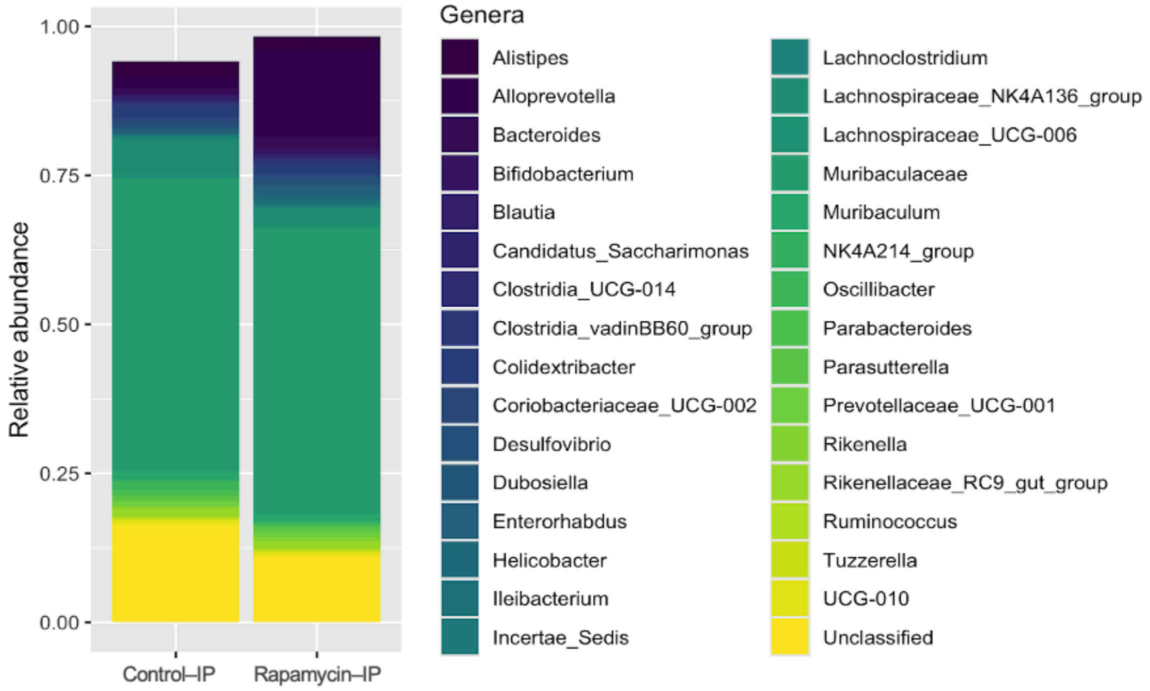


**Fig. S6 Genus composition**
Han2021 (IP); stacked bars show mean relative abundance per group (Control–IP, Rapamycin–IP); colours denote genera; inclusion thresholds: detection 1.25% and prevalence ≥50%; “Other” pools low-abundance genera and “Unclassified” denotes reads not assigned at genus level; the profile was dominated by Muribaculaceae, *Alistipes*, and *Bacteroides*, with a notable Unclassified fraction; genus-level composition did not differ between groups (Kruskal–Wallis p = 0.6193)

#### Firmicutes-to-Bacteroidetes Ratio

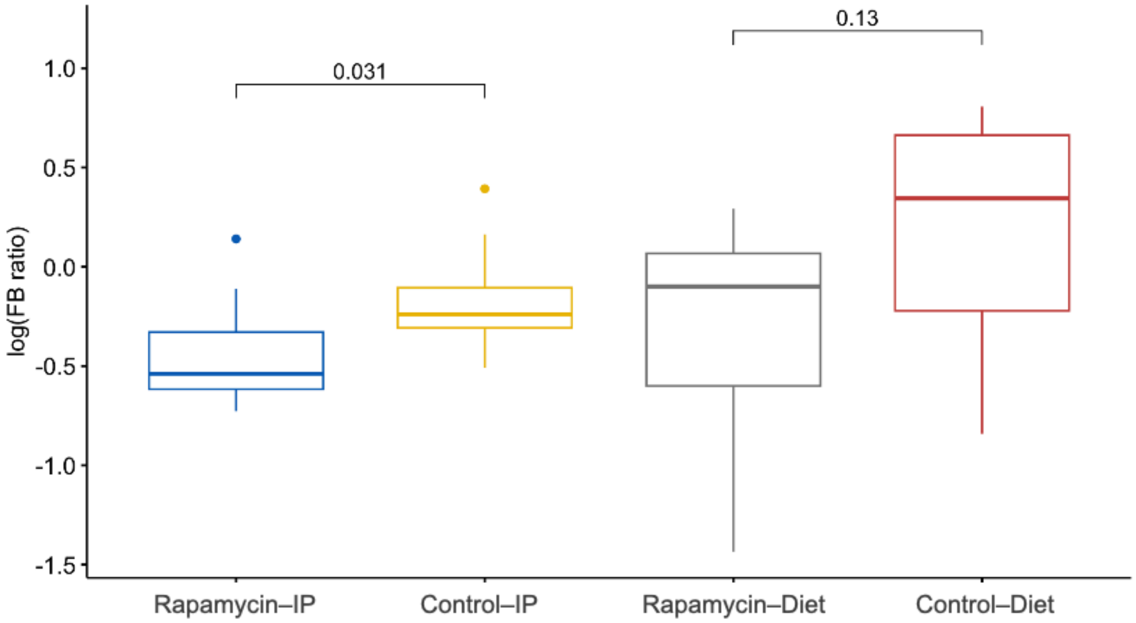


**Fig. S7 Log-transformed Firmicutes-to-Bacteroidota (F/B) ratio — Bitto2016 (IP and Diet)**
Boxplots show the median and interquartile range with whiskers extending to 1.5×IQR and outliers as points; bracket labels are two-sided Wilcoxon rank-sum p-values; comparisons: Rapamycin–IP vs Vehicle–IP p = 0.031 and Rapamycin–Diet vs Control–Diet p = 0.13; values are ln(Firmicutes/Bacteroidota), where lower values indicate a lower F/B ratio

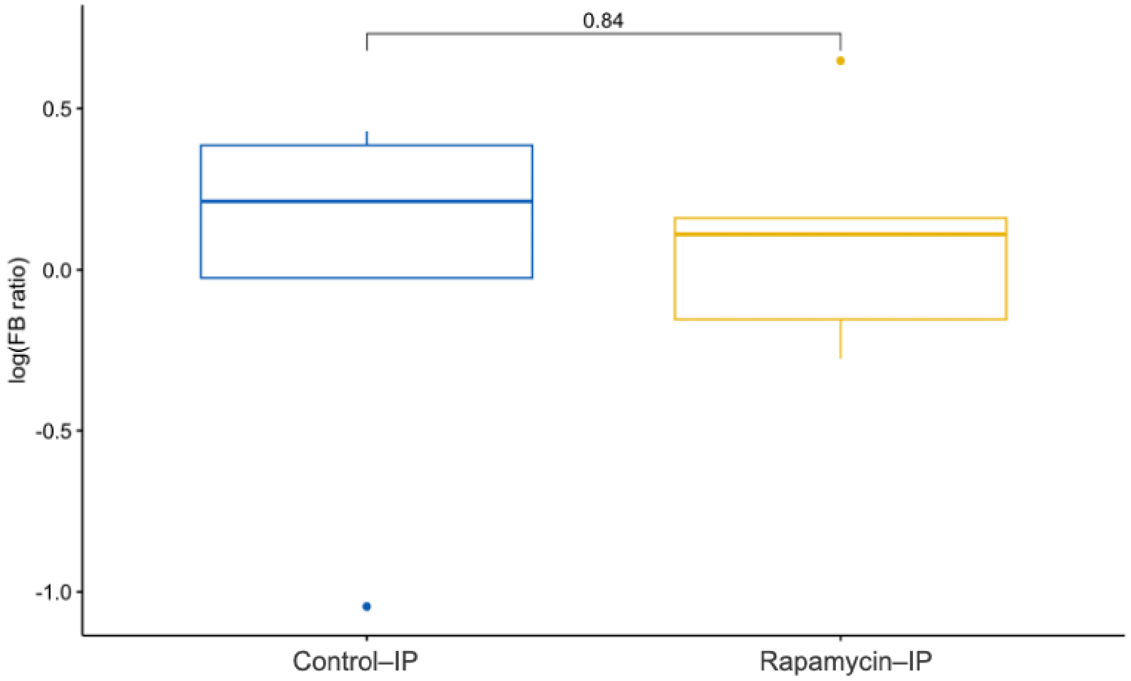


**Fig. S8 Log-transformed Firmicutes-to-Bacteroidota (F/B) ratio — Yang2022 (IP)**
Boxplots show the median and interquartile range with whiskers extending to 1.5×IQR and outliers as points; the bracket label gives the two-sided Wilcoxon rank-sum p-value for Control–IP vs Rapamycin–IP (p = 0.84); values are ln(Firmicutes/Bacteroidota), where lower values indicate a lower F/B ratio

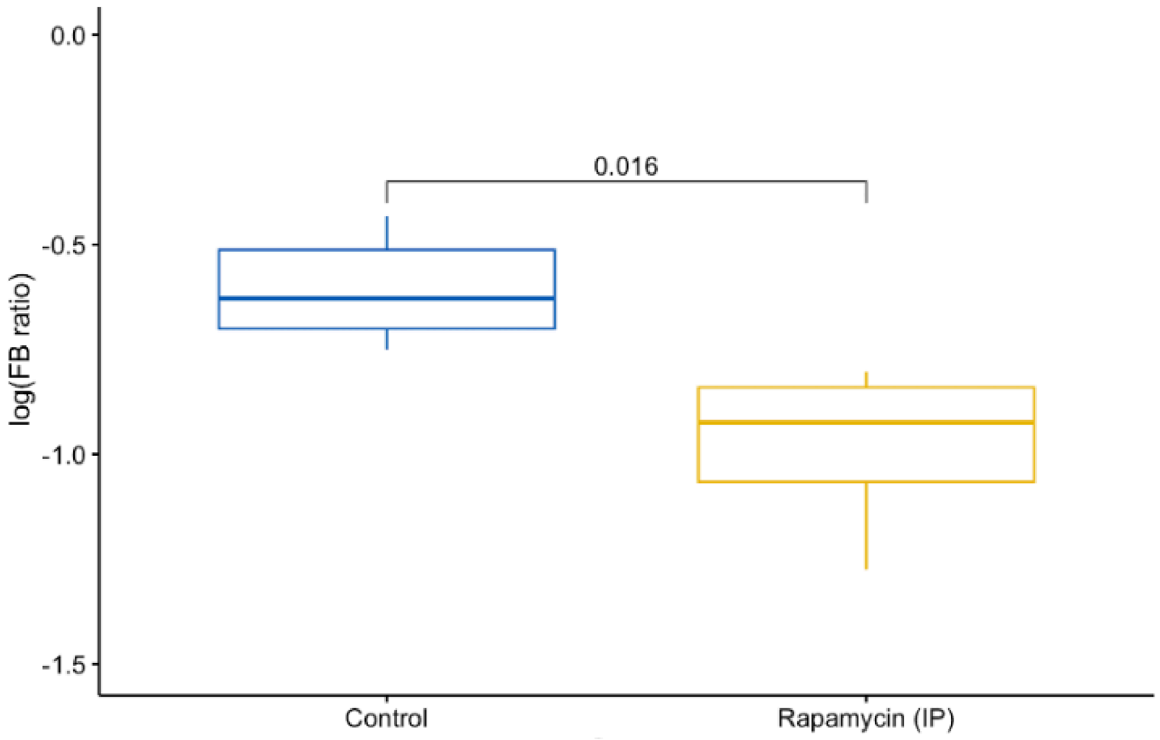


**Fig. S9 Log-transformed Firmicutes-to-Bacteroidota (F/B) ratio — Han2021 (IP)**
Boxplots show the median and interquartile range with whiskers extending to 1.5×IQR and outliers as points; the bracket label gives the two-sided Wilcoxon rank-sum p-value for Control–IP vs Rapamycin–IP ($p = 0.016$); values are ln(Firmicutes/Bacteroidota), where lower values indicate a lower F/B ratio

#### Core Microbiome Analysis

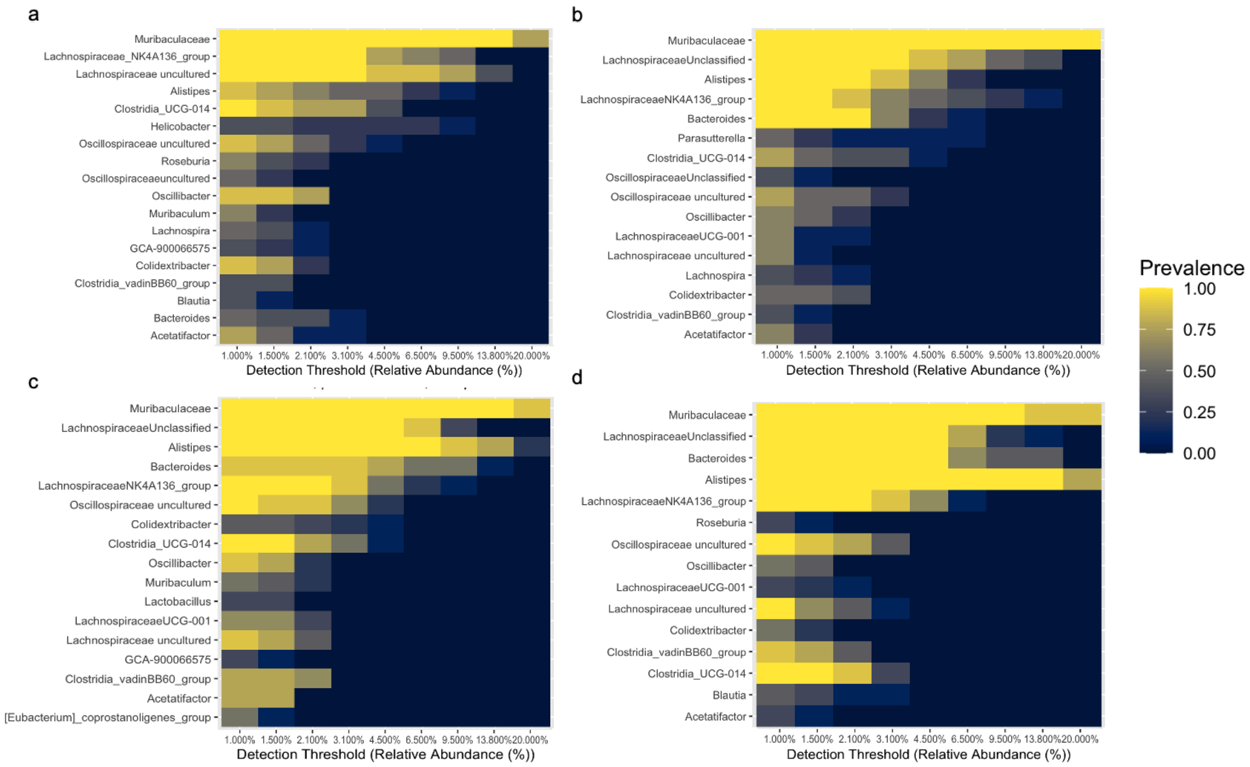


**Fig. S10 Core genus-level microbiome composition across treatment groups in Bitto2016**
Heatmaps show genus prevalence across increasing detection thresholds of relative abundance (x-axis, %). Core membership was defined as detection ≥0.1% with prevalence ≥80%; only genera with minimum prevalence ≥30% in any group are shown. Panels: (a) Control–IP, (b) Rapamycin–IP, (c) Control–Diet, (d) Rapamycin–Diet. The core community was largely conserved across groups, with Muribaculaceae, Lachnospiraceae NK4A136 group, *Alistipes*, and *Bacteroides* recurring in all panels.

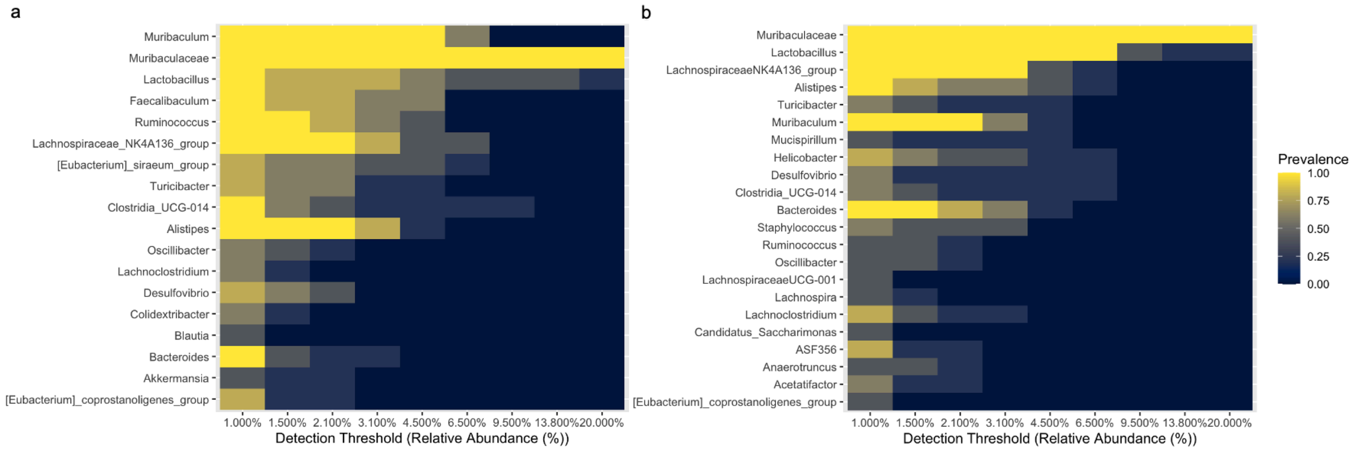


**Fig. S11 Core genus-level microbiome composition in Yang2022 dataset**

Heatmaps show genus prevalence across increasing detection thresholds of relative abundance (x-axis, %). Core membership was defined as detection ≥0.1% with prevalence ≥80%; only genera with minimum prevalence ≥30% in either group are shown. Panels: (a) Control–IP, (b) Rapamycin–IP. Shared core taxa included Muribaculaceae, Lachnospiraceae NK4A136 group, and *Alistipes*, while *Faecalibaculum* and *Lactobacillus* were prominent in controls but reduced with rapamycin

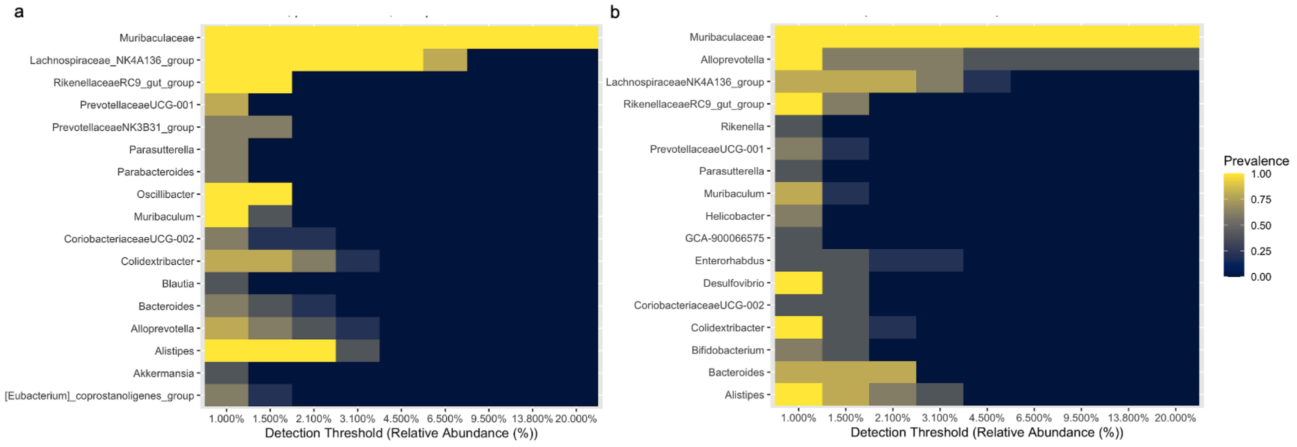


**Fig. S12 Core genus-level microbiome composition in Han2021 dataset**

Heatmaps show genus prevalence across increasing detection thresholds of relative abundance (x-axis, %). Core membership was defined as detection ≥0.1% with prevalence ≥80%; only genera with minimum prevalence ≥30% in either group are shown. Panels: (a) Control–IP, (b) Rapamycin–IP. *Alistipes* was more prevalent in controls along with higher representation of the Lachnospiraceae NK4A136 group, whereas the rapamycin group showed increased core prevalence of *Alloprevotella* and *Bacteroides*; Muribaculaceae remained dominant in both groups.

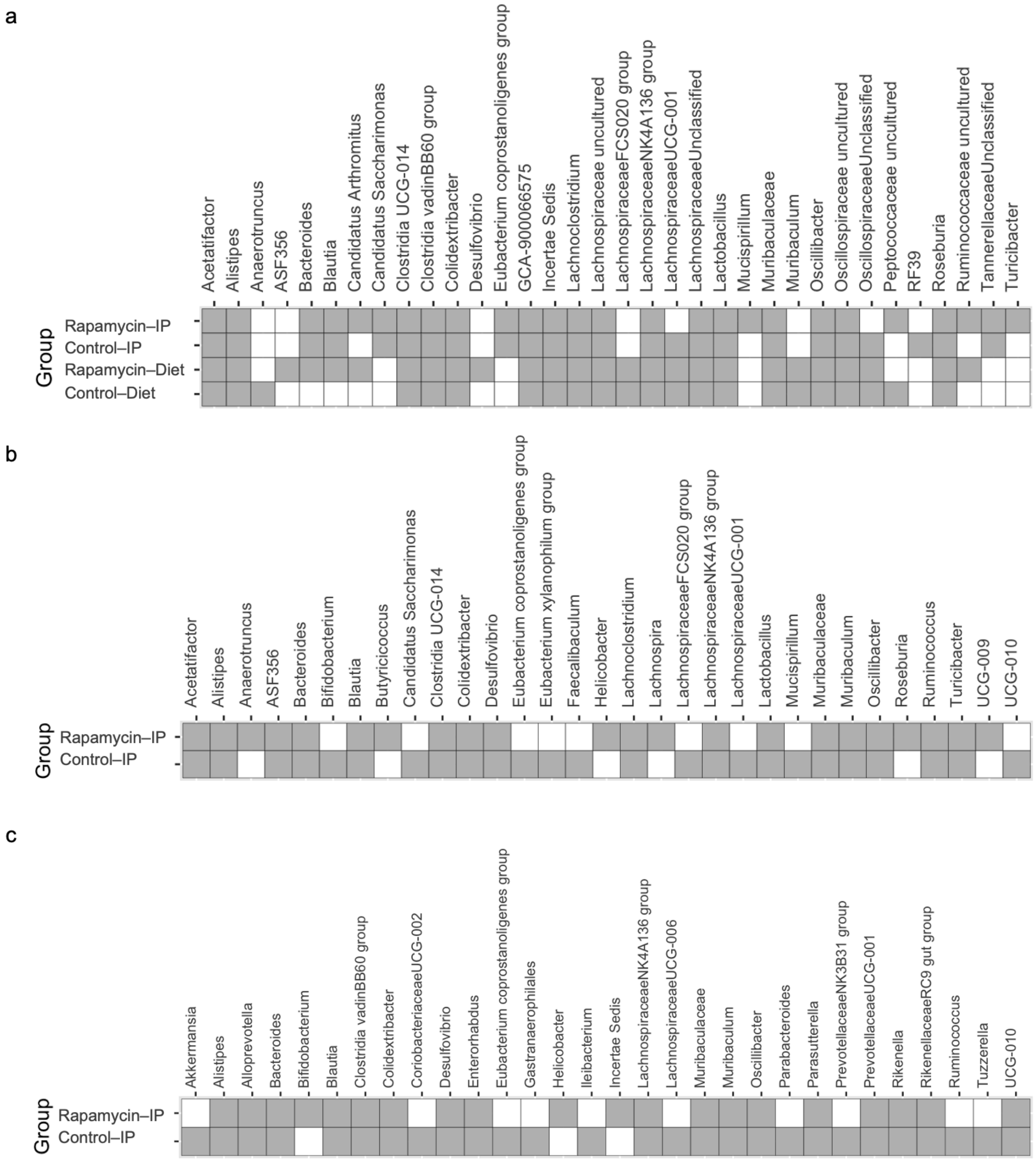


**Fig. S13 Core genus presence across treatment groups**

Presence/absence matrices depict bacterial genera meeting core microbiome criteria (detection threshold ≥0.1% relative abundance and prevalence ≥80%) across three datasets: (a) Bitto 2016 (Control intraperitoneal (IP), Rapamycin–IP, Rapamycin–Diet, Control–Diet, (b) Yang 2022 (Control–IP, Rapamycin–IP, and (c) Han 2021 (Control–IP, Rapamycin–IP. Grey tiles indicate genera detected in the core microbiome for the respective group; white tiles indicate absence. Muribaculaceae and Lachnospiraceae NK4A136 group were consistently observed as core members, regardless of treatment. *Helicobacter* showed increased core prevalence in rapamycin-treated groups in two datasets. In contrast, genera such as *Eubacterium* spp. and *Blautia* displayed variable responses to treatment across studies.

#### Diversity Analysis

##### Alpha Diversity

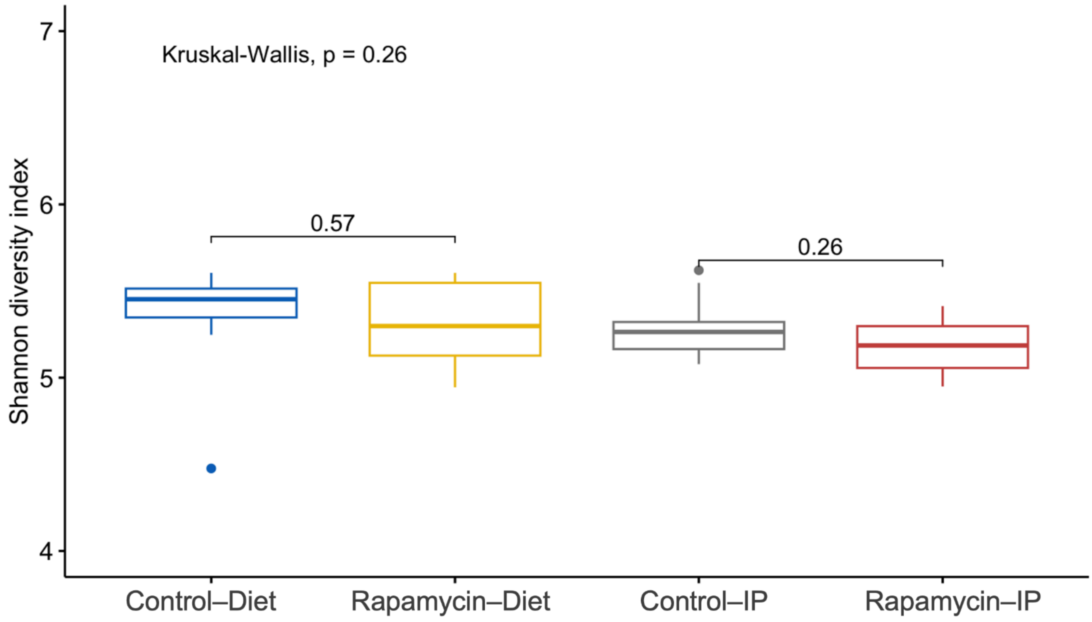


**Fig. S14 Shannon diversity — Bitto2016 (IP and Diet)**
Boxplots show the median and interquartile range with whiskers to 1.5×IQR and outliers as points; overall differences across the four groups were not significant (Kruskal–Wallis p = 0.26); Rapamycin–Diet showed a slight upward shift versus Control–Diet (Wilcoxon p = 0.57) and Rapamycin–IP a slight downward shift versus Control–IP (Wilcoxon p = 0.26); taken together, rapamycin produced no consistent increase or decrease in Shannon diversity in this dataset

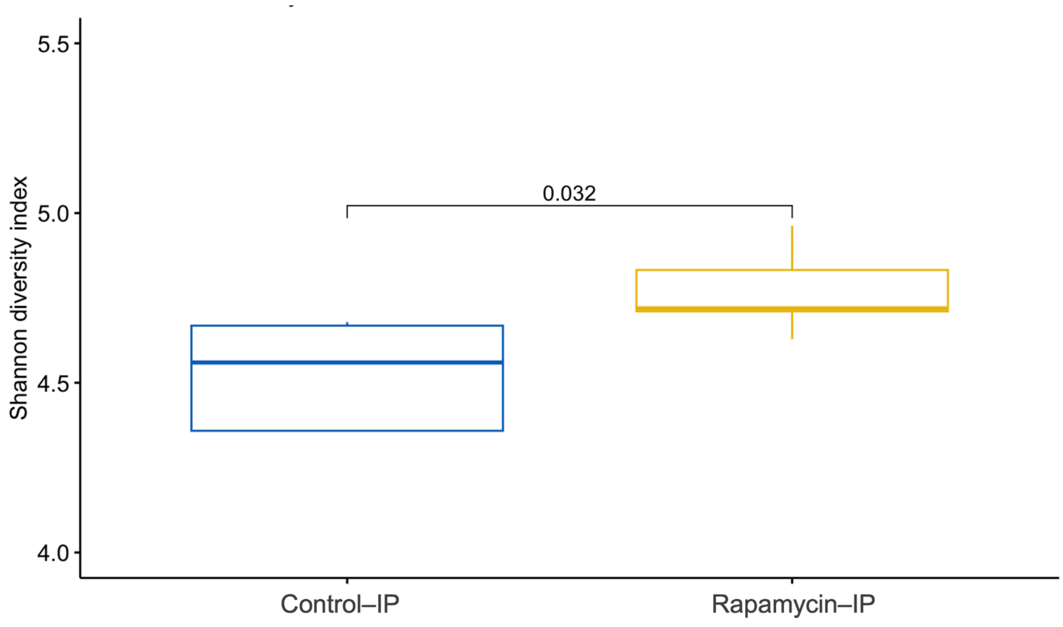


**Fig. S15 Shannon diversity — Yang2022 (IP)**
Boxplots show the median and interquartile range, and whiskers to 1.5×IQR with outliers as points. The Rapamycin–IP group shows an upward shift relative to Control–IP; groups compared with a two-sided Wilcoxon rank-sum test, Control–IP vs Rapamycin–IP p = 0.032

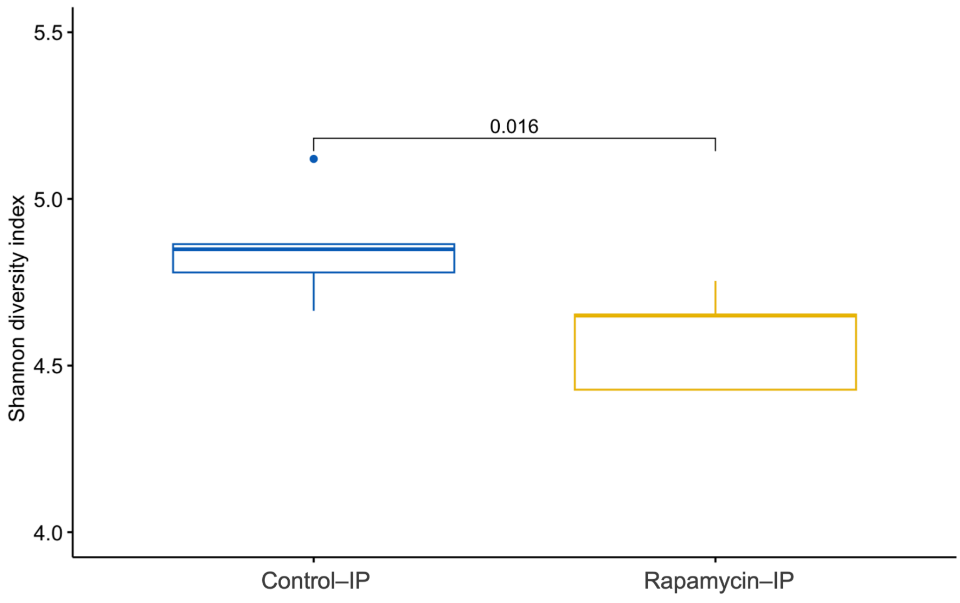


**Fig. S16 Shannon diversity — Han2021 (IP)**
Boxplots show the median and interquartile range with whiskers to 1.5×IQR and outliers as points; Rapamycin–IP showed a downward shift versus Control–IP (two-sided Wilcoxon p = 0.016), indicating lower alpha diversity in the rapamycin group

#### Beta Diversity

##### Weighted UniFrac Analysis

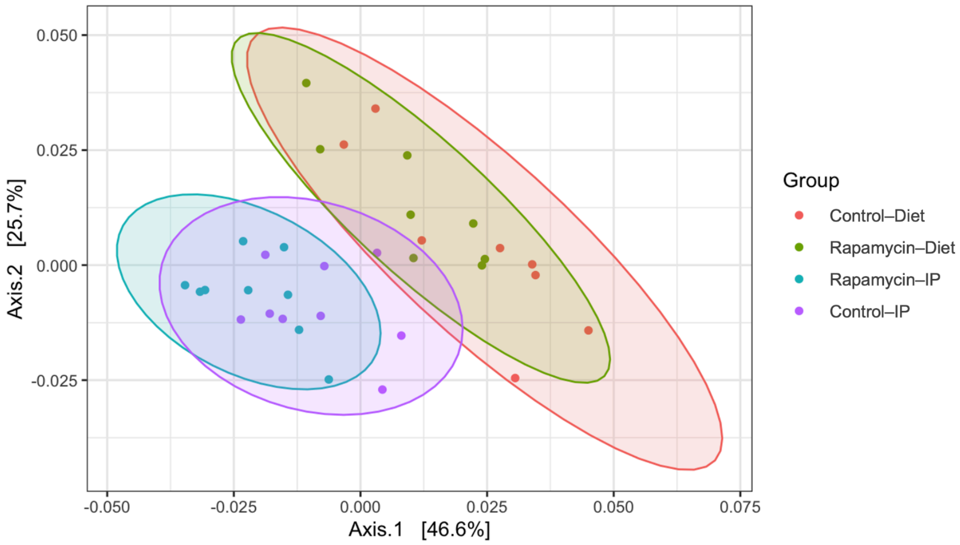


**Fig. S17 PCoA of weighted UniFrac distances — Bitto2016**
Points are coloured by group (Control–Diet, Rapamycin–Diet, Rapamycin–IP, Control–IP); ellipses depict normal-approximation group dispersion; Axis 1 and Axis 2 explain 46.6% and 25.7% of variance, respectively; overall differences by PERMANOVA/adonis across groups: p = 0.0001 (df = 3); dispersion differed among groups (betadisper ANOVA F = 3.32, p = 0.033; permutation p = 0.041), so separation reflects both centroid location and variability

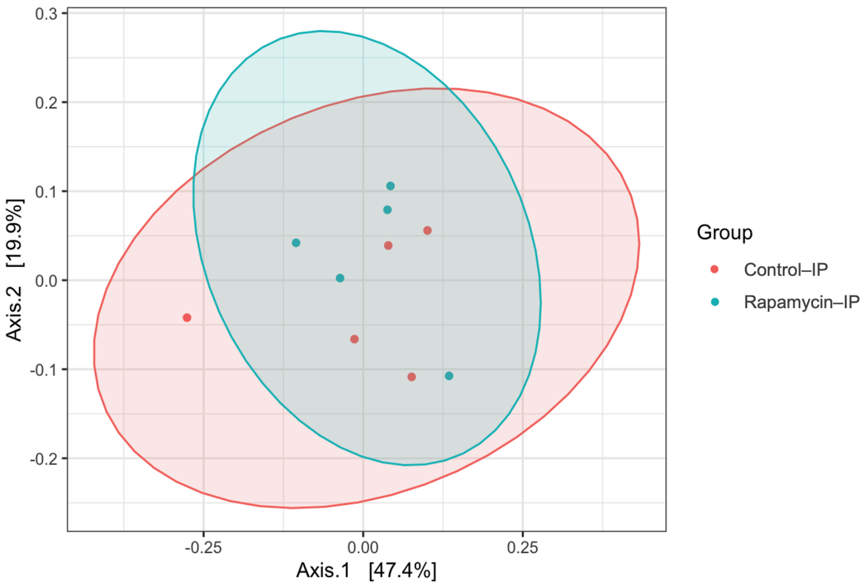


**Fig. S18 PCoA of weighted UniFrac distances — Yang2022 (IP)**
Points are coloured by group (Control–IP, Rapamycin–IP) with normal-approximation ellipses showing group dispersion; Axis 1 and Axis 2 explain 47.4% and 19.9% of variation, respectively; overall group differences were not significant by PERMANOVA/adonis (p = 0.39, df = 1), and dispersion did not differ between groups (betadisper permutation p = 0.63); the substantial overlap in the ordination indicates no detectable treatment effect on community structure

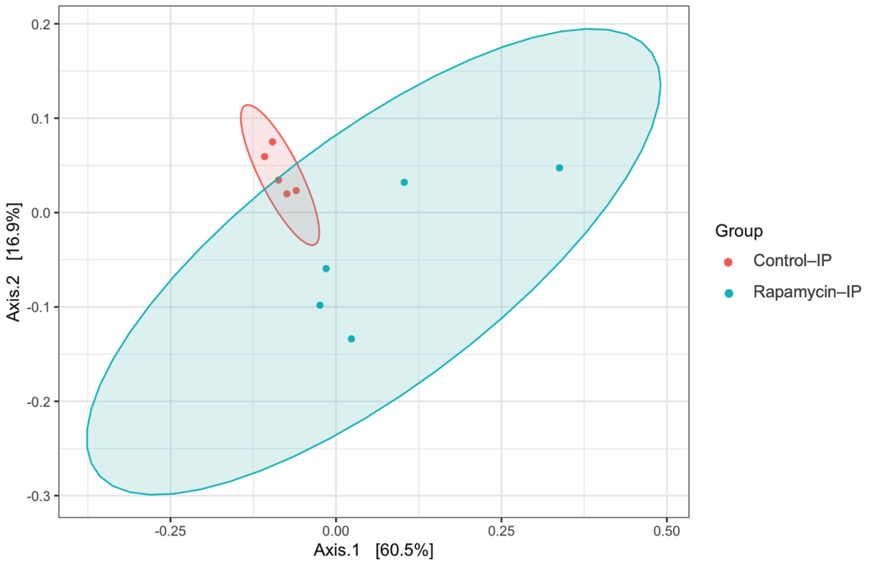


**Fig. S19 PCoA of weighted UniFrac distances — Han2021 (IP)**
Points are coloured by group (Control IP, Rapamycin IP) and ellipses show normal-approximation group dispersion; Axis 1 and Axis 2 explain 60.5% and 16.9% of the variation; overall group differences were significant by PERMANOVA/adonis (p = 0.0004; df = 1); the dispersion test was also significant (betadisper permutation p = 0.022), indicating unequal within-group dispersion

#### Unweighted UniFrac Analysis

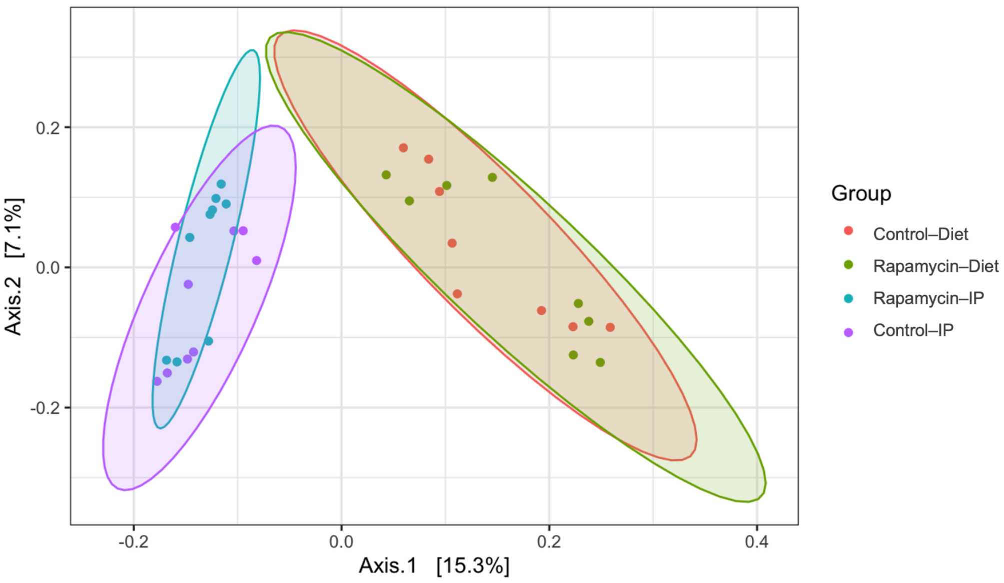


**Fig. S20 PCoA of unweighted UniFrac distances — Bitto2016**
Points are coloured by group and ellipses show normal-approximation dispersion; Axis 1 and Axis 2 explain 15.3% and 7.1% of the variation; samples given diet cluster on the right of Axis 1 (Control–Diet, Rapamycin–Diet) and samples given IP cluster on the left (Control–IP, Rapamycin–IP); within each route the control and rapamycin clouds largely intermingle, while the diet groups—especially Rapamycin–Diet—show a broader spread than the IP groups; overall differences across the four groups were significant by PERMANOVA/adonis (p = 0.0001; df = 3); the homogeneity-of-dispersion test was also significant (betadisper permutation p = 0.001), indicating unequal within-group spread

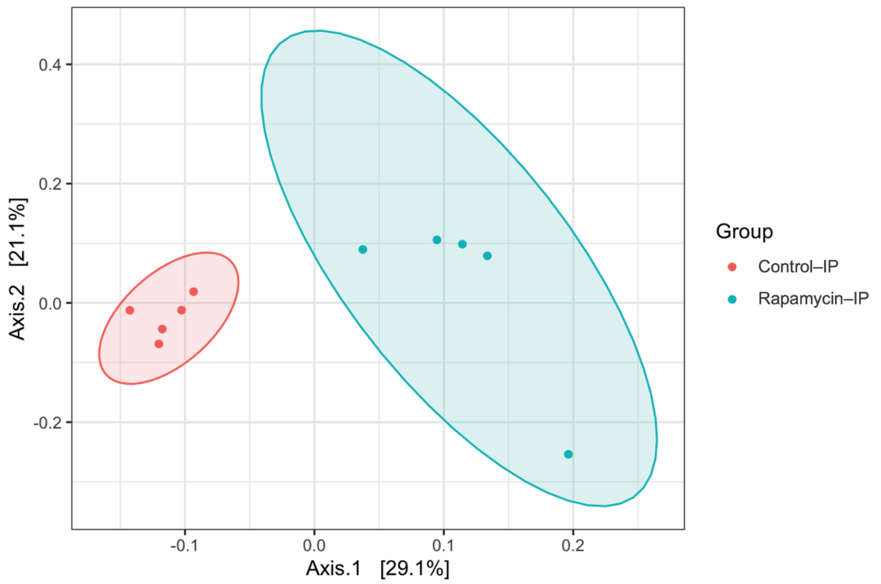


**Fig. S21 PCoA of unweighted UniFrac distances — Yang2022 (IP)**
Points are coloured by group (Control IP, Rapamycin IP) and ellipses show normal-approximation group dispersion; Axis 1 and Axis 2 explain 29.1% and 21.1% of the variation; the test for homogeneity of multivariate dispersion was significant (permutation p = 0.039), indicating unequal within-group spread; overall group differences in centroids were not significant by PERMANOVA/adonis

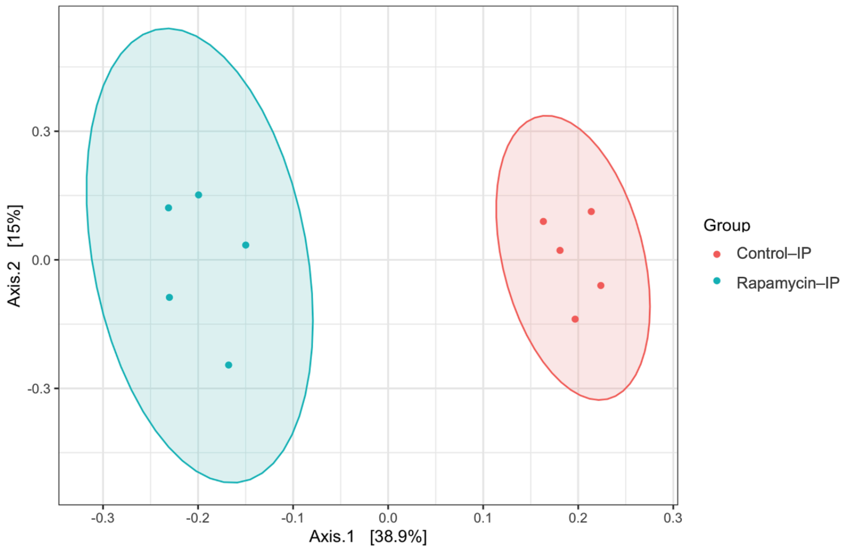


**Fig. S22 PCoA of unweighted UniFrac distances — Han2021 (IP)**
Points are coloured by group (Control–IP, Rapamycin–IP) and ellipses show normal-approximation group dispersion; Axis 1 and Axis 2 explain 38.9% and 15.0% of the variation; overall group differences were significant by PERMANOVA/adonis (p = 0.0074; df = 1); dispersion also differed by permutation test (p = 0.043)

#### Bray-Curtis Dissimilarity Analysis

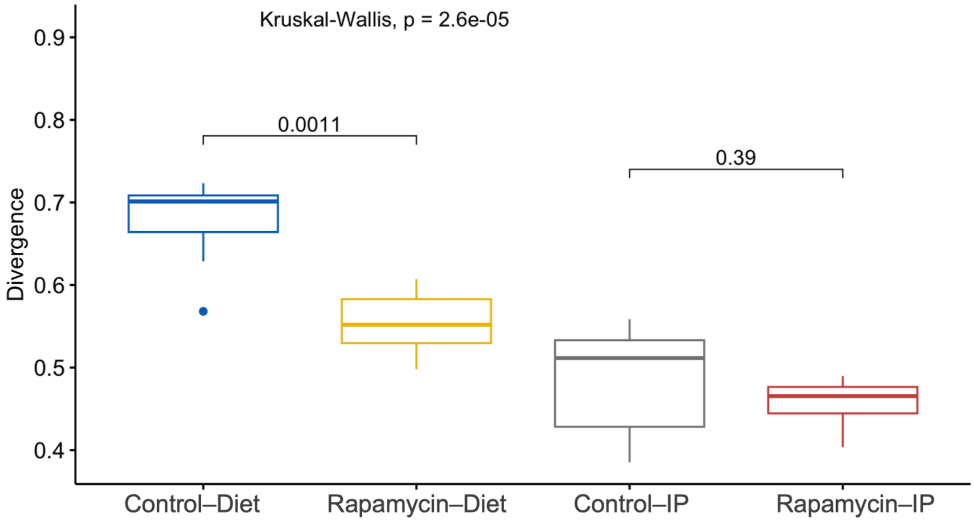


**Fig. S33 Bray–Curtis divergence in Bitto2016**
Boxplots show distance to each group's average community in PCoA space (median, IQR, whiskers to 1.5×IQR; outliers as points). Overall group differences were significant (Kruskal–Wallis $p = 2.6\times10^{-5}$). Control–Diet showed higher divergence than Rapamycin–Diet (Wilcoxon p = 0.0011), while Control–IP vs Rapamycin–IP was not significant (p = 0.39). Food groups tended to be more divergent than IP groups.

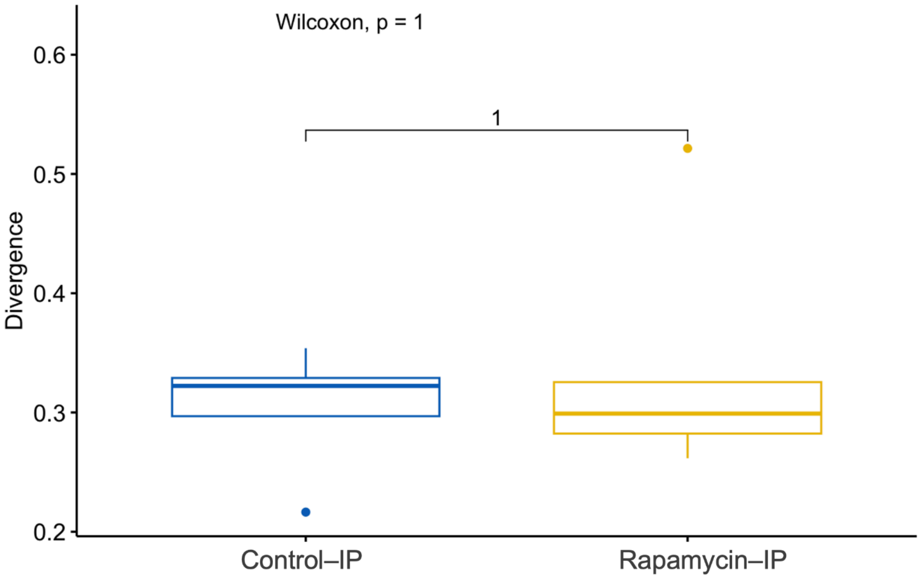


**Fig. S24 Bray–Curtis divergence (Yang2022 – IP)**
Boxplots show each sample's distance to its group average community (median, IQR, whiskers to 1.5×IQR; outliers as points). Control–IP and Rapamycin–IP were indistinguishable (two-sided Wilcoxon rank-sum p = 1.00), indicating similar inter-individual divergence in both groups.

#### Differential Abundance analysis

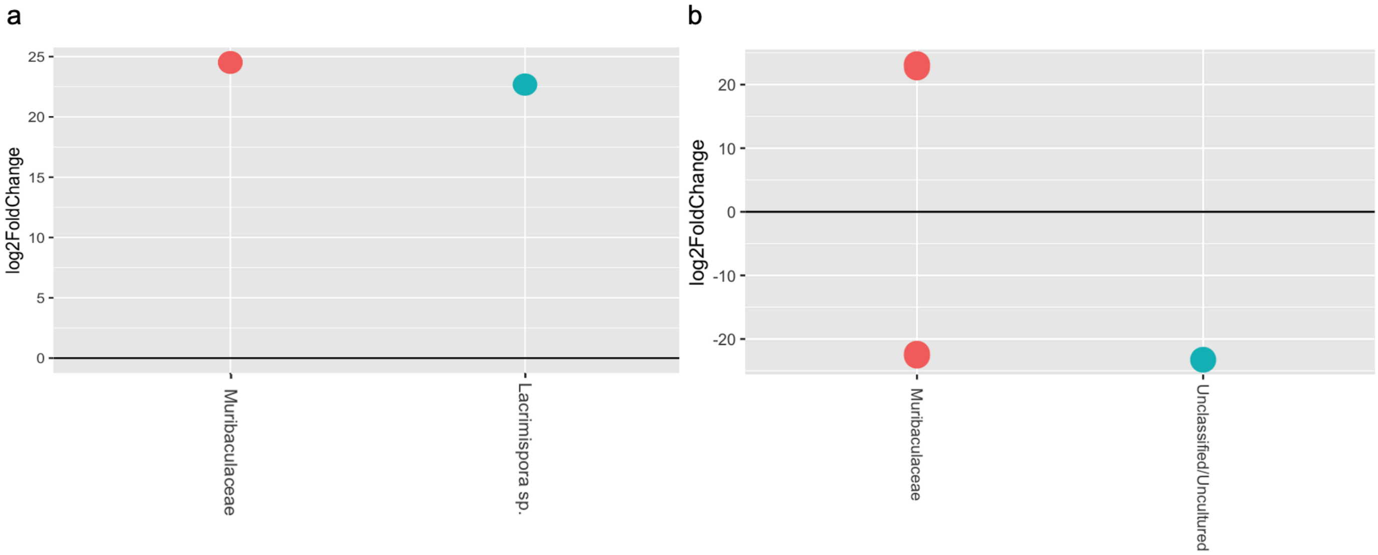


**Fig. S25 Differentially abundant taxa in Bitto2016**

DESeq2 contrasts within route (Wald test; Benjamini–Hochberg FDR < 0.05); points show $Log_2FC$ for rapamycin vs control (a) Rapamycin–IP vs Vehicle–IP: Muribaculaceae ($Log_2FC > +23$) and *Lacrimispora* sp. ($Log_2FC > +20$) enriched; (b) Rapamycin–Diet vs Control–Diet: Muribaculaceae ASVs showed bidirectional responses (some $Log_2FC > +20$, others $< -20$) and Unclassified/Uncultured were depleted ($Log_2FC < -25$); very large $|Log_2FC|$ values typically reflect near-zero abundance in one group and should be interpreted primarily for direction rather than exact magnitude

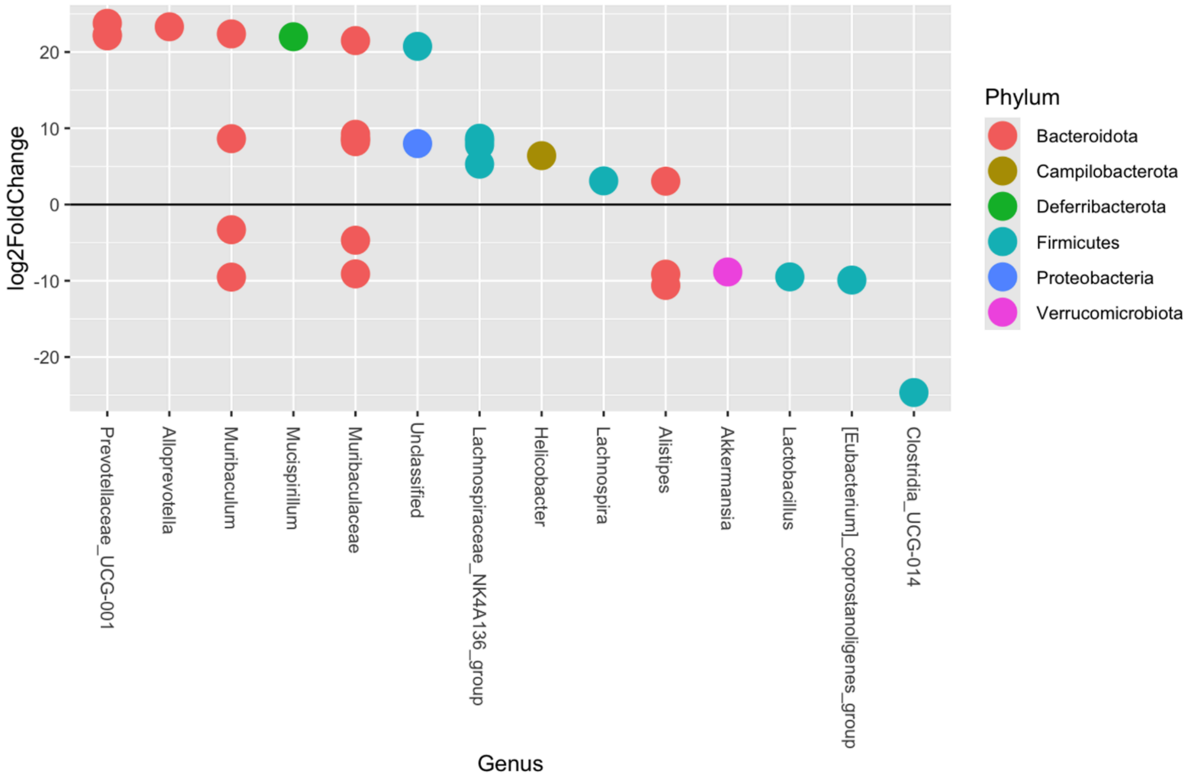


**Fig. S26 Differentially abundant genera in Yang2022**

DESeq2 contrast Rapamycin–IP vs Control–IP (Wald test; BH FDR < 0.05; detection ≥0.25%); points show $\log_2$ fold change and colours indicate phyla. Increases were observed for Prevotellaceae *UCG-001, Alloprevotella, Mucispirillum (*Deferribacterota), *Helicobacter,* and several Firmicutes including *Lachnospira* and the Lachnospiraceae NK4A136 group. *Muribaculum* and Muribaculaceae showed mixed behaviour with both enriched and depleted ASVs, consistent with strain-level heterogeneity. Decreases included *Akkermansia, Lactobacillus, [Eubacterium] coprostanoligenes* group, and *Clostridia* UCG-014, which showed the largest negative effect

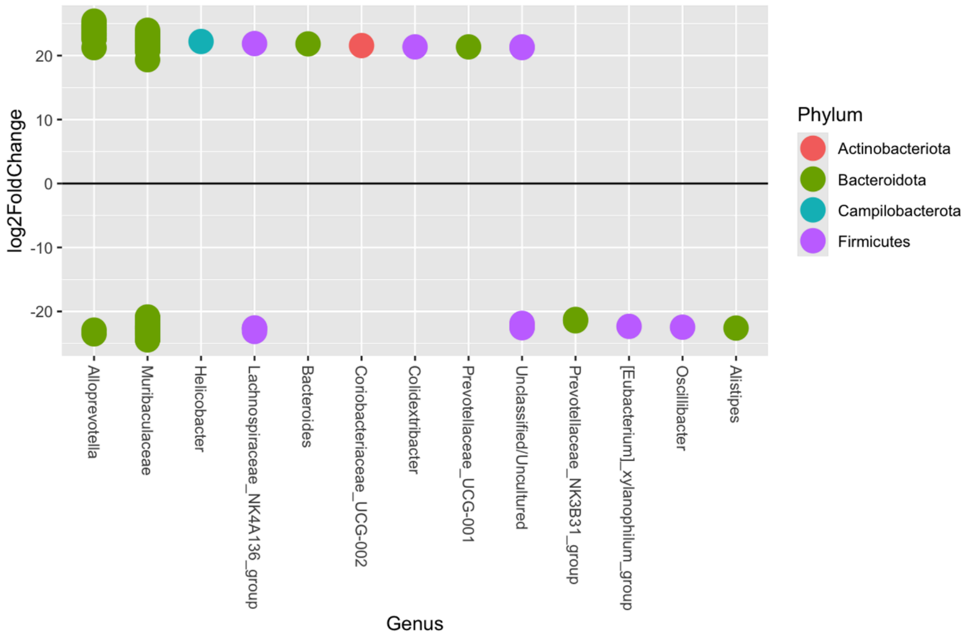


**Fig. S27 Differentially abundant genera in Han2021**

DESeq2 contrasts Rapamycin–IP vs Control–IP (Wald test; BH FDR < 0.05; detection ≥0.25%); points show $\log_2$ fold change and colours indicate phyla. Enriched under rapamycin were *Helicobacter, Bacteroides,* Coriobacteriaceae *UCG-002, Colidextribacter,* Prevotellaceae UCG-001 and a subset of Muribaculaceae ASVs. Depletions included *Alloprevotella, Alistipes, Oscillibacter, [Eubacterium] xylanophilum* group, Prevotellaceae NK3B31 group, Lachnospiraceae NK4A136 group, Unclassified/Uncultured and additional Muribaculaceae ASVs. The opposing directions within Muribaculaceae indicate ASV-level heterogeneity

#### Functional Pathway Analysis

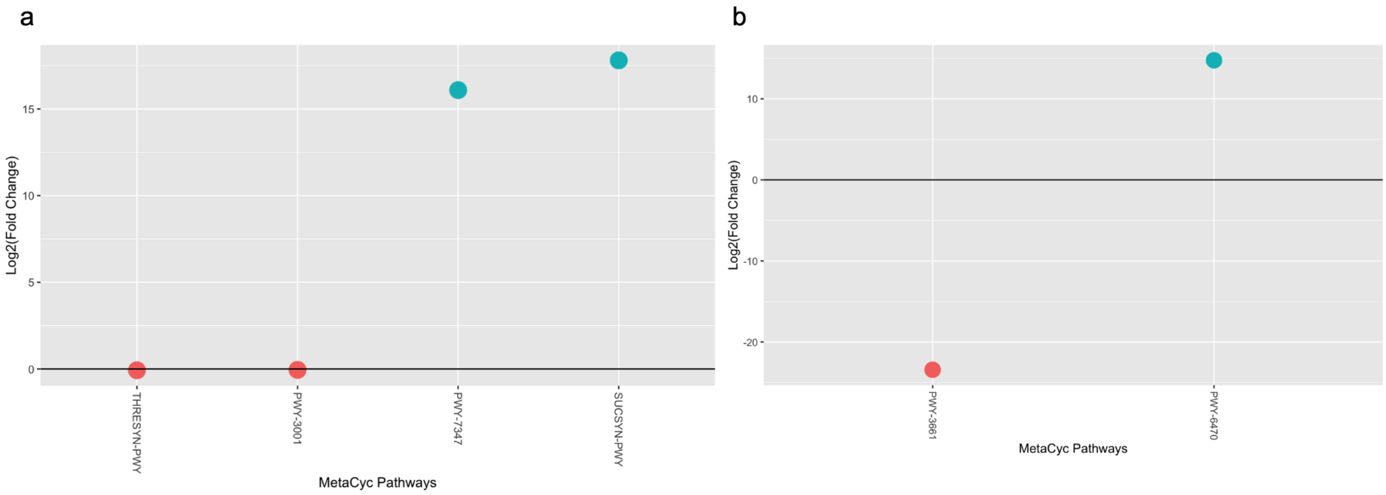


**Fig. S28 Predicted functional pathways in Bitto2016**

Differential pathway abundance from PICRUSt2 + DESeq2 contrasts within route (Rapamycin vs Control; Wald test with Benjamini–Hochberg correction); only pathways with FDR < 0.05 are shown, points are $\log_2$ fold change relative to control with 0 marking no difference

(a) Intraperitoneal: Sucrose biosynthesis I (SUCSYN-PWY) and Sucrose biosynthesis III (PWY-7347) increased strongly (both >10 $\log_2$FC), while the Superpathway of L-threonine biosynthesis (THRESYN-PWY) and L-isoleucine biosynthesis I from threonine (PWY-3001) decreased modestly (b) Dietary: Glycine betaine degradation II (PWY-3661) decreased and Peptidoglycan biosynthesis V (β-lactam resistance) (PWY-6470) increased, indicating changes in osmotic stress handling and cell-wall remodelling

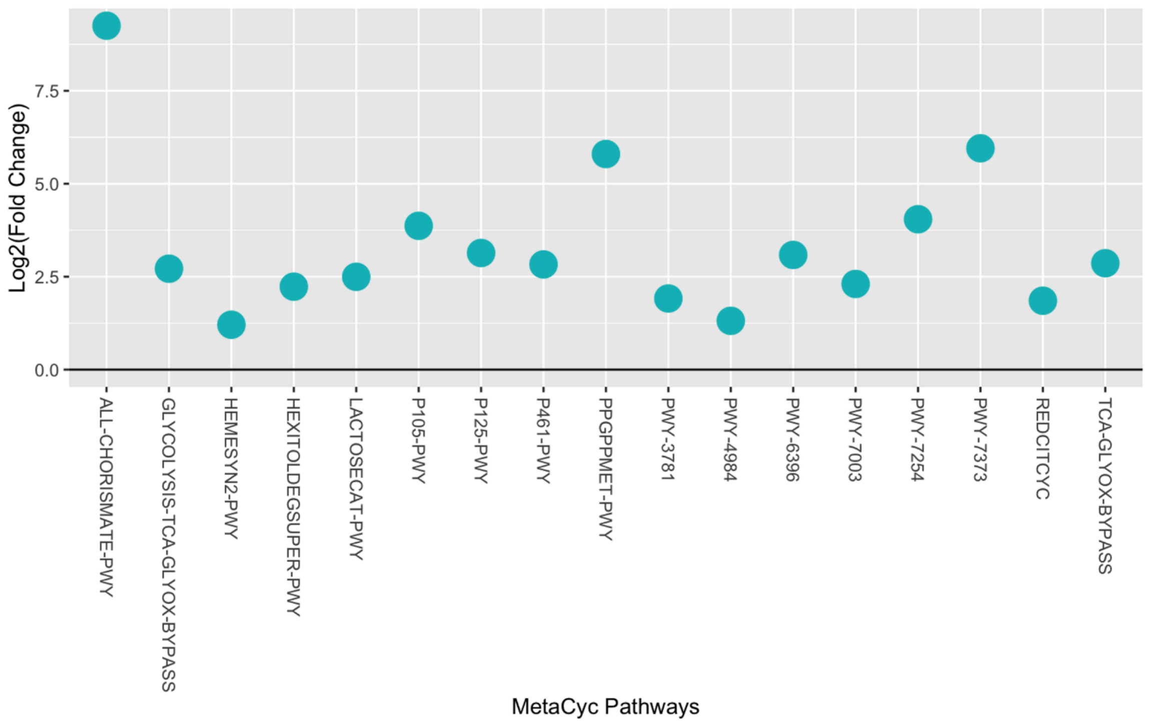


**Fig. S29 Predicted functional pathways in Yang2022**

Differential pathway abundance for Rapamycin–IP vs Control–IP (PICRUSt2 + DESeq2, Wald test with Benjamini–Hochberg correction); only pathways with FDR < 0.05 are shown, points are $log_2$ fold change relative to control with 0 marking no difference

Significant enrichments included Chorismate biosynthesis (ALL-CHORISMATE-PWY), ppGpp stringent-response (PPGPPMET-PWY), Demethylmenaquinone biosynthesis (PWY-7373), TCA/glyoxylate-bypass variants (PWY-7254, REDCITCYC, TCA-GLYOX-BYPASS, GLYCOLYSIS-TCA-GLYOX-BYPASS), 2,3-butanediol biosynthesis (P125-PWY, PWY-6396), Lactose degradation (LACTOSECAT-PWY), Hexitol degradation superpathway (HEXITOLDEGSUPER-PWY), and Oxygen-independent heme biosynthesis (HEMESYN2-PWY), consistent with increased aromatic-precursor supply, stress-response activation, and re-routing of central carbon metabolism under IP rapamycin
